# *De novo* design of transmembrane accessory subunits for fold stabilization and expansion

**DOI:** 10.64898/2026.05.14.725059

**Authors:** Sebastian Jojoa-Cruz, Saba Kanwal, Nolan P. Jacob, Weiyi Tang, Mio Murakoso, Minghao Zhang, Jiayi Li, Masy Domecillo, Nicholas F. Polizzi, John R. Yates, Huong T. Kratochvil, Fabio P. Gomes, Heedeok Hong, Marco Mravic

**Affiliations:** Department of Integrative Structural and Computational Biology, The Scripps Research Institute, La Jolla, CA, 92037, USA; Department of Chemistry, Michigan State University, East Lansing, MI 48824, USA; Department of Chemistry, University of North Carolina at Chapel Hill, Chapel Hill, NC 27599, USA; Department of Cancer Biology, Dana-Farber Cancer Institute, Boston, MA, USA; Department of Biological Chemistry and Molecular Pharmacology, Harvard Medical School, Boston, MA, USA; Department of Chemistry, The Scripps Research Institute, La Jolla, CA, 92037, USA; Department of Chemistry, Virginia Commonwealth University, Richmond, Virginia 23284, USA; Department of Biochemistry & Molecular Biology, Michigan State University, East Lansing, MI 48824, USA

## Abstract

Transmembrane (TM) proteins play essential roles in biology as transporters, ion channels, chaperones, enzymes, and mediators of signal transduction. However, membrane proteins often suffer from inefficient folding and intrinsic instability. Misfolding in cells can cause numerous loss-of-function pathologies. Likewise, denaturation upon purification in the laboratory is a critical barrier to structure determination and characterization of key biochemical mechanisms. Generalizable strategies to stabilize membrane proteins remain limited. Here, we developed an informatics-based *de novo* design strategy to create synthetic auxiliary subunits that interact with the TM helices of a model pentameric ion channel, thereby bolstering folding while maintaining channel function. Biochemical and structural characterization reveal the synthetic TM subunits can also be used to create larger multi-spanning designer proteins of custom topology. This proof-of-concept motivates the feasibility of computationally designed accessory TM helices as potential pharmacological chaperone “folding correctors” of membrane proteins in disease and as tools in structural biology.

## INTRODUCTION

Folding and assembly of transmembrane (TM) domains define the structures and stabilities of membrane proteins. Mutations and other chemical changes that disrupt interactions between TM spans often result in reduced protein stability, misfolding, and aberrant cellular trafficking, which contribute to loss-of-function and pathological mechanisms driving disease^1,2^. Using exogenous molecules to correct folding and trafficking mechanisms is a promising therapeutic approach for many membrane protein classes^3^. FDA-approved Trikafta drugs for cystic fibrosis exemplify this strategy, boosting functional CFTR ion channel levels by stabilizing vulnerable folding intermediates^4,5^. However, methods for generating new chemical matter aimed at stabilizing integral membrane protein folding are far less evolved than those established for water-soluble proteins, with the lipid bilayer posing distinct challenges to molecular discovery and design.

In Nature, transmembrane accessory subunits are a prevalent mechanism by which protein complexes are stabilized and their functions diversified within membranes. Some TM adaptors exert chaperone-like effects, such as boosting stability or redirecting membrane trafficking, while others tune efficacy, selectivity, and downstream cellular activity. Notable examples are TM adaptors of GPCRs (*e.g*., RAMP family^6,7^, CD69^8^) and ion channel auxiliary subunits like the KCNE family, which modify voltage-gated potassium channels^9^ (K_v_’s). Here, we explore the feasibility and utility of using protein design to create analogous synthetic accessory subunits for membrane fold stabilization (**Fig. 1a**). As well, we test if the *de novo* TM adaptor domains are a reliable approach to expand simple protein assemblies into larger more complex multi-spanning folds of desired topologies (**Fig. 1b**).

**Figure 1.**
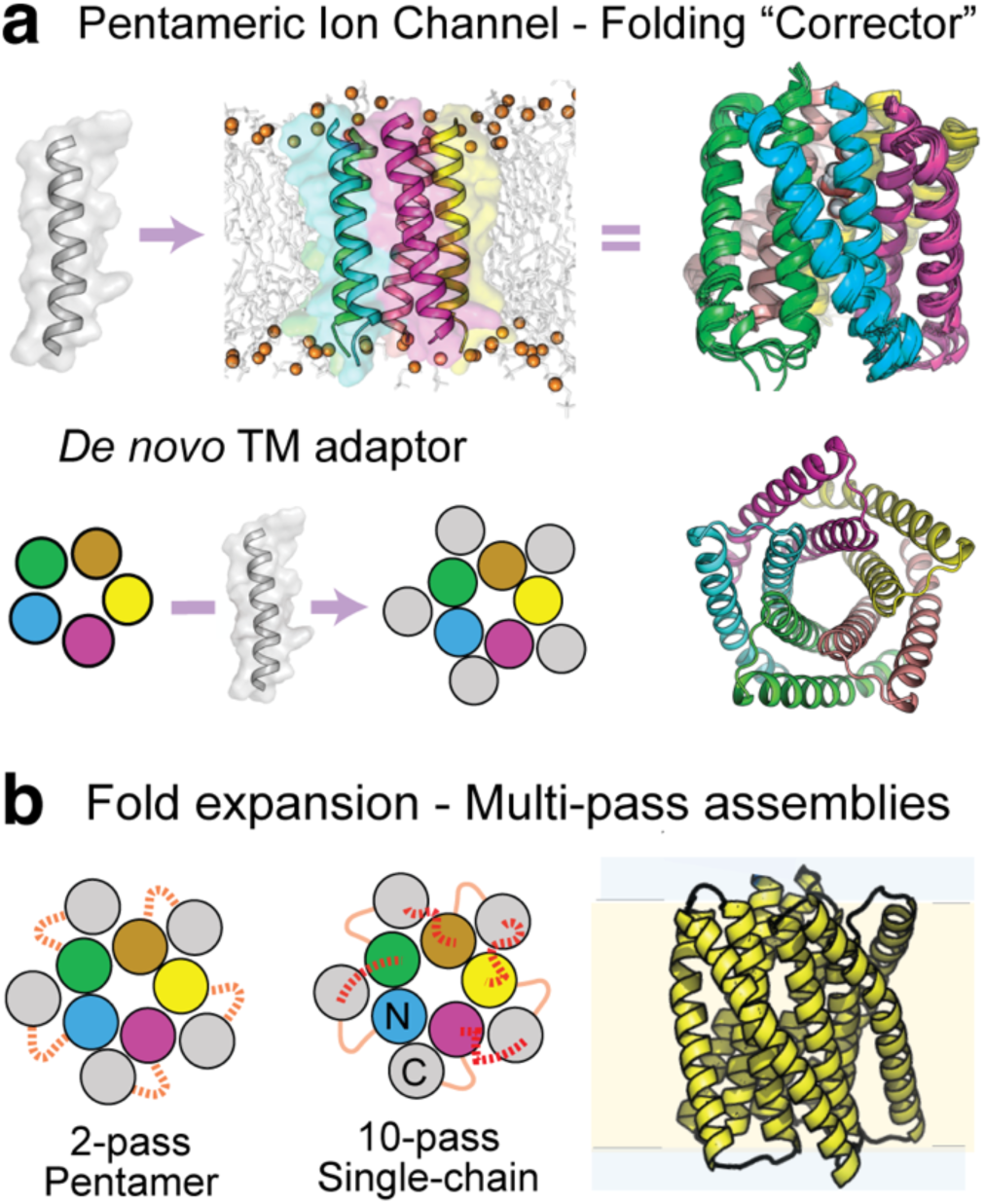
Design of transmembrane fold stabilizing domains. (**a**) Left, de novo accessory subunits (grey) to buttress assembly of a single-span pentameric ion channel “LQLL” (colors). Right, fused protein of multi-pass subunits. **(b)** Design of large complex synthetic membrane proteins by fusing TM structural building blocks via de novo loops (orange or red, solid and dashed lines).

Most *de novo* designs of multi-pass helical membrane proteins of defined structure thus far have been re-engineered from known stable water-soluble analogs^10–13^ rather than building sequences from the ground up to fold withiin lipid environments. As well, new engineered interaction complexes with membrane proteins have mostly been limited to those between single-spanning TM helices (*e.g.*, of single-pass receptors)^14^, except for very recent tethered GPCR adaptors^15^. The membrane design field would benefit from approaches to create new interacting polypeptides which can complement and target the dynamic and often kinked, irregular complex multi-helical surfaces present in most protein families (transporters, channels, etc). The ability to engineer accessory complexes and stabilize more membrane protein classes would be broadly enabling, with potential to enable new functions in synthetic biology or to facilitate therapeutic stabilization of misfolding or mistrafficking diseases^4,16,17^.

As a proof-of-concept, we debut a structural informatics strategy to design TM adaptors interacting with a pentameric ion channel, then biochemically characterize the extent to which the synthetic TM domains stabilize its folding. We targeted variants of this ion channel with tendency to disassemble or misfold to lower non-functional stoichiometries. A lead *de novo* accessory TM subunit resulted in channel assemblies with improved folding specificity and stability (*i.e*., to proteases, thermal denaturation, short-chain detergents) with the intended structure observed by cryo-electron microscopy (cryo-EM). The many synthetic stable protein assemblies demonstrate that this TM adaptor design strategy is a versatile approach for correction and synthetic expansion of membrane protein folds.

## RESULTS

### Design of *de novo* TM accessory subunits

Recent studies described a family of synthetic homo-pentameric ion channels assembled from single-span TM peptides which achieve high proton selectivity by recruiting hydration within a predominantly apolar central pore^18^. The most conductive ion channel variant, named LQLL, bearing a central glutamine within the pore, is primarily pentameric in SDS-PAGE. However, when its oligomeric distribution is observed in mild detergents, separation by size-exclusion chromatography (SEC) reveal that LQLL adopts a mixture of monomers and dimers alongside a primary higher-order oligomer, the likely conductive pentameric assembly (**Fig. 2a**). Over time and upon incubation in short-chain detergents, the LQLL protein disassembles to a greater portion of non-conducting dimers and monomer (**Fig. 2a**). The instability and lower folding specificity of this ion channel appears analogous to misfolding and disassembly for many natural membrane proteins in disease^19^. The most stable but non-conducting variant, named “eVgL” (or “LLLL”) adopts a pentamer which is remarkably resistant to denaturation (SDS, Urea, 95° C)^20^. However, despite this stability, pentameric folding is incomplete; a small fraction of monomer always persists in mild detergents (**Fig. 2a**). We hypothesized that new interactions from a designed accessory TM domain could further promote folding, leaving negligible non-pentameric fraction and yielding a more biochemically uniform sample. Thus, we used these proteins and their behaviors as controlled model systems for testing the hypothesis that synthetic TM subunits can be made on-demand with effects akin to those of pharmacological chaperone “folding correctors” of ion channels and GPCRs^4,21^.

**Figure 2.**
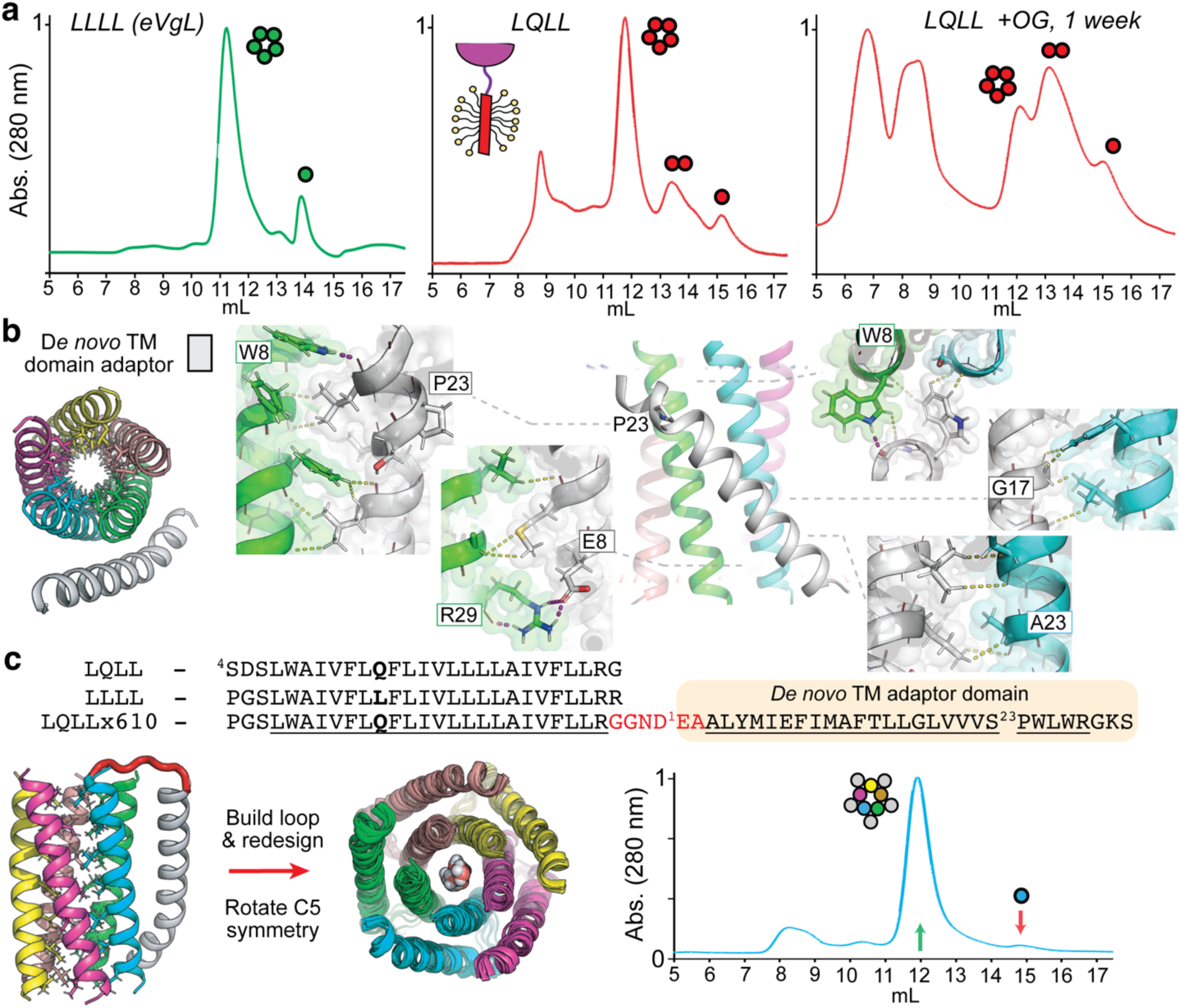
Folding and stabilization of designed pentameric ion channel variants. (**a**) Size exclusion chromatography (SEC) UV chromatograms of purified SUMO fusion proteins (cartoon) of TM domains LLLL-eVgL (green; left) and the less stable ion channel variant LQLL (red; middle, directly after purification; right, after 1 week incubated in 200 mM octyl-glucoside) in a 3 mM myristyl sulfobetaine (C14B) aqueous mobile phase, s200i column, representatives of n=3 experiments. Primary sequences listed; pore-facing mutation bolded. Colored circles denote oligomeric state assigned to SEC peaks. **(b)** Left, design of *de novo* TM adaptor (gray) to pentameric assembly inter-subunit interface. Right, interaction layers within top design TM610, optimizing packing apolar sidechains with close backbone contacts (<3.5 Å, yellow dash) and inter-helical polar interactions (purple dash; H-bonds, salt bridges). Conserved proline, sticks. **(c)** Top, TM sequences of LQLL proton channel, LLLL-eVgL non-channel, and new 2-pass LQLLx610 (loop, red; *de novo* designed accessory TM domain, orange, expected TM spans, underlined). Lower left, model of LLLL pentamer (color, cartoon; apolar core as sidechain sticks) and bound TM adaptor helix (gray) with synthetic loop built (red). Middle, overlayed MD simulation frames of 2-pass pentamer LQLLx610, with water in the pore. Right, SEC trace of 2 mg/mL SUMO-LQLLx610pass in 3 mM C14B mobile phase (s200i column), denoting increase in oligomeric fraction (green arrow) and reduced unfolded monomer peak to negligible fraction (red arrow); representative of n=3 experiments.

We proposed a strategy of binding a synthetic TM domain at the pentamer’s membrane-exposed inter-subunit interface (**Fig. 1a, 2b**). Adding conformationally specific intermolecular interactions should reinforce intra-pentamer packing and folding energetics. This design of new interactions complementary to an existing membrane protein fold requires molecular recognition akin to binder design. However, successful strategies of model building and sequence design in membranes for this design task to date have been largely limited to interactions between isolated helices, using monomeric binding single-spanning TM helices to target single-span receptors^22–25^. Thus, for this more complex objective, we devised a knowledge-based model building approach that precisely positions new helical binding elements conditioned on the specific geometry of the target: here, the pentamer’s inter-subunit groove (**Fig 2b, S1**). Accessory helix backbone templates are drawn from known, structurally similar membrane helix packing structures in nature^26,27^, thus are compatible to host stabilizing sequences within these tertiary geometries. This strategy circumvents the broad, mostly unproductive sampling of helix conformations by stochastic docking^28^ and parametric methods^10,29^. This data-directed approach provides distinct helix binding geometries compared to those generated from diffusion-based^30^ methods, which enrich for rigid ideal α-helices.

Our method conducts a two-stage search, first querying a database of natural membrane protein structures for instances of TM helices geometrically similar to the targeted binding surface: LQLL’s two parallel-oriented inter-subunit interface (15-residue helix-helix fragments) (**Fig S1a**). Secondly, we sub-search the local environment around each membrane protein structure where the queried target helix-helix instance is found (<2.0 Å RMSD), seeking to identify additional TM domains nearby to the targeted helix groove (**Fig. S1b**). These resulting data-mined additional helices can then be structurally aligned to the targeted protein and subsequently used as backbone templates for design of compatible interacting sequences.

This first-stage search identified many instances of the ion channel’s inter-subunit helix-helix geometry in diverse natural proteins (**Fig. S1b**), unsurprisingly, since it is a known common TM building block of straight ideal α-helices^20,26^. Sub-searching protein structures that surround instances of helix geometries similar to LQLL’s 2-helices (i.e., the search “bait”) reveals that a high frequency of those instances, 38%, have a third compatibly interacting TM domain nearby (**Fig S1a-b**). Analyzing hundreds of these data-mined three-helix arrangements identifies a strong geometric preference amongst “prey” TM helices^27^, with 64% interacting antiparallel relative to the target’s mutually parallel “bait” helices. This data guided us to focus on antiparallel backbone templates for design.

Structural clustering of >100 antiparallel “prey” TM helices which contact both “bait” TM helices identified a dominant binding mode that varied modestly across clusters (**Fig. S1b**) hosting a highly conserved proline-induced kink (**Fig. 2b, S2a**). The kink facilitates the single helix to bridge interactions with both LQLL TM subunits in a continuous interface lining the pentamer’s tilted inter-subunit groove. While helix distortions are seldom incorporated in protein design deliberately, they are common in natural membrane proteins to support conformational flexibility, structural diversity, and precise sidechain positioning^31^. We therefore used templates containing this shared kink, expecting intermolecular interactions surrounding the designed proline will encode the desired accessory helix geometry and compensate intramolecular distortion.

Upon analyzing the data-mined natural TM helices structurally aligned to the pentamer groove, we deduced that this knowledge-based strategy identifies and positions helical elements pre-disposed for sidechain knobs-in-holes (KIH) packing (**Fig. S1c**). For interface residues within the positioned helix templates, sidechain vectors are frequently directed towards the “holes” (i.e. the inter-residue gap that exposes polar mainchain area to lipid) on the targeted pentamer helices. For example, one such natural “prey” helix, TM4 of GPCR S1PR1^32^, when positioned into LQLL’s inter-subunit groove forms compatible KIH packing at several residue clusters without any prior sequence optimization at its C-terminal half (**Fig. S1c**). Its N-terminal half is more distant, but sidechains still project towards potential holes on both LQLL subunits, well-suited for redesign to optimize new interactions. Thus, our model building approach navigates explicit data-driven preferences for sequence-structure guidelines and yields “design-able”^26^ helical templates, i.e. backbone arrangements able to accommodate many distinct sequences and favorable KIH packing.

Next, sequence design of TM accessory helices in complex with the pentamer was conducting using rotamer trials with the implicit RosettaMembrane model^33^ from the top 2 helix template clusters (**Fig. S2a-b**). We biased residue selections based on natural amino acid preferences amongst the “prey” TM helices data-mined (**Fig. S2a-b**). The recurring internal proline was fixed to stabilize the helix kink, as was a threonine at the preceding position.

In model evaluation, a key design principle was to optimize van der Waals (vdW) interactions meeting a specific geometric criterion we hypothesize as critical for apolar packing to confer stability amongst competing lipid tails within the membrane environment. That is, sidechain KIH packing events interdigitating tightly into the adjacent helix backbone: <3.2 Å to mainchain heavy atoms (**Fig. 2b**). This backbone-directed vdW geometry is enriched in many high stability TM interfaces and has demonstrated promise as a structural quality metric in membrane protein design^20,23,24,34^. Top designs were filtered by surface complementarity, number of inter-helical hydrogen bonds, and interface computed energy, then ranked based on the number of backbone-directed packing events. Combined with sequence clustering and all-atom molecular dynamics simulations of ∼50 models for stability in model membranes (**Fig. S2b-d**), these criteria led to a final set of 11 designs, all sharing 2 keystone polar interactions (**Fig. 2b**). Near their N-terminus, each design coordinates LQLL’s Arg28 either by salt bridge via an acidic residue or by H-bond via a serine or threonine. Near their C-terminus, designs all accept an intermolecular H-bond from LQLL’s Trp8 amide at the backbone carbonyl group preceding (i-2) the conserved proline, precisely positioned to facilitate and stabilize the kinked conformation.

Although this protein design was completed prior to release of AlphaFold (AF), we found 5 of 11 designs were confidently predicted by AF3 to adopt the intended pentamer binding mode (**Fig. S3, Table S2**). The proline kink is recapitulated alongside key KIH and polar interactions. Thus, this approach yields high-quality TM domain sequences and structures with potential to stabilize the pentameric ion channel.

### Interaction and fold stabilization of *de novo* TM accessory domains

To investigate whether the synthetic accessory TM domains in isolation could bind the target pentamer, we used cell-based and *in vitro* experiments as cursory screens for lead candidates. Using the NanoBit-based “memPPI” split enzyme protein-protein interaction assay^35^, we tested the binding propensity of each designed TM domain for the eVgL pentamer upon co-expression in HEK293T cells (**Table S3**). Across designs, expression-normalized interaction propensities were similar to those of non-interacting negative controls (**Fig. S4**). Given many cellular factors could shroud detection of TM complexes of modest strength, such as co-localization, trafficking, topology, and relative binder:target molar ratios, we did not rule out possibility of a complex. Instead, we performed an *in vitro* screen experiment. Fluorophore-labeled TM peptides (∼4 kDa) of each design were mixed with the pentamer (as a sumo-fusion protein, ∼100 kDa oligomer), then monitored their ability to co-elute in a binding complex in SEC. Upon mixing peptide with pentamer at ratios between 0.2 to 1.2 molar equivalents in the relatively harsh mistryl-sulfobetaine (C14B) at detergent concentrations well below crowding (>= 1 protein per micelle), we observed minor fractions of designed TM peptides co-eluting with the high molecular weight pentamer (**Fig. S5**), indicating binding to the assembly as expected. The top design, TM610, has the largest co-eluting peak amongst those tested (**Fig. S5b-c**). Although a complex was detected, its low occupancy hinders clear characterization and interpretation of how the accessory helix’s binding may modulate the pentamer folding.

To facilitate testing of whether a designed TM accessory subunit can stabilize the misfolding-prone LQLL ion channel, we fused the TM610 to the C-terminus of the channel via a short flexible linker. The resulting two-pass construct (LQLLx610), in which the second TM span is the *de novo* “folding corrector” subunit, was purified and characterized (**Fig. 2c, Table S4**). By SEC, LQLLx610 folds to a single oligomeric state of pentamer size in all detergents tested (C14B, DDM, C8E4) with or without an N-terminal sumo fusion protein (**Fig. 2c, S6**), as do pore-modifying variants (i.e. LNLLx610). Monomers and non-conducting smaller oligomers become negligible (>98% fraction folded). Upon challenge to disassembly by extended incubation in short-chain detergent (200 mM OG), this dominant LQLLx610 folded oligomer remains intact as the dominant species (>95%) even months later. Thus, folding behavior with the de novo “folding corrector” subunit is improving from that of the parent LQLL-1pass channel, and for drastically longer time spans (**Fig. S6c**).

The 2-TM span proteins can spontaneously refold both in octyl-glucoside and in SDS after purification in organic solvent (**Fig 3a, S6h**). They run on SDS-PAGE as higher molecular weight oligomers, which can be denatured by heat (95*°* C), but migrate anomalously fast (∼25 kDa observed versus 35 kDa expected). This behavior is consistent with a folded TM assembly in SDS micelles^28,36^ and matches migration of the parent 1-pass pentameric channels^18,20^. Glutaraldehyde cross-linking of SUMO-LQLLx610 results in patterns of protein oligomers up to 5 subunits (**Fig 3b**), confirming folding to the pentameric assembly as expected.

**Figure 3.**
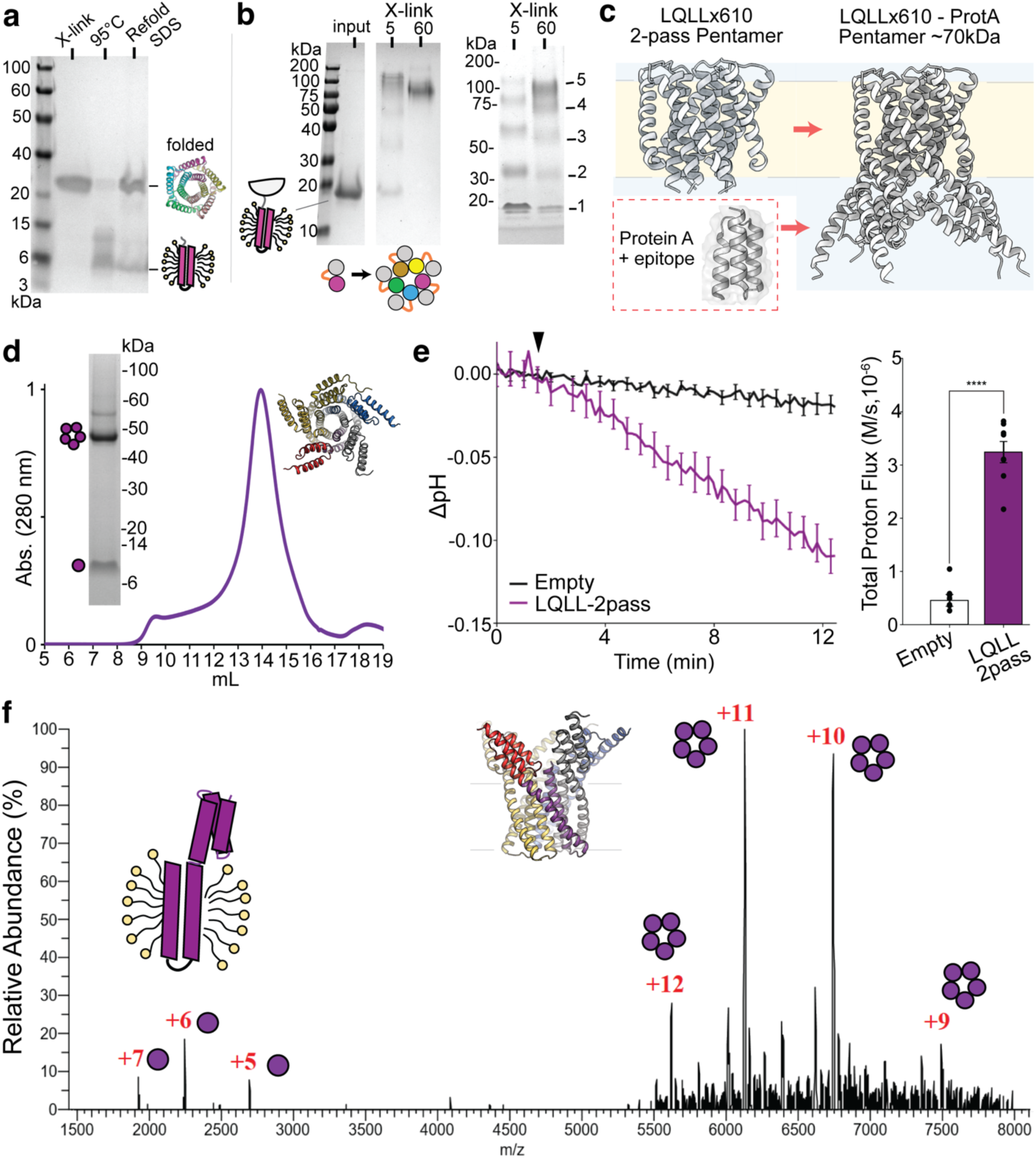
Pentameric assembly and stability of LQLLx610 proteins. (**a**) SDS-PAGE of cleaved re-folded LQLLx610 (7 kDa monomer) after purification in organic solvent. Lane 1, reconstituted to 0.5 mg/mL in 50 mM octyl-glucoside and cross-linked (“X-link) by glutaraldehyde. Lane 2 OG-reconstituted protein heated to 95 °C. Lane 3 reconstituted in 2% w/v SDS (lane 3) leading to an SDS-resistant oligomer in Tris-Glycine-SDS gel. **(b)** SDS-PAGE of SUMO-LQLLx610 (20 kDa monomer) before and after glutaraldehyde cross-linking (“X-link). Protein bands migrate with different behavior in Tris-Glycine-SDS (left) or MES-SDS (right) gel running system, but reveal folding of 2-pass subunits to a pentameric stoichiometry (cartoon; helices, colored circles) **(c)** Fold expansion by rigid helix fusion of 3-helix bundle protein A domain to LQLLx610 C-terminus. **(d)** SEC UV trace of cleaved LQLLx610-proteinA fusion (13.8 kDa monomer) in 1 mM DDM, runs as single major monodisperse folded species. Inset, SDS-resistant oligomer by gel electrophoresis, also migrating anomalously (Expected: ∼70 kDa 10-pass assembly). **(e)** Proton conduction in liposome flux assay for empty liposomes (no protein, black) and liposomes with cleaved LQLLx610-proteinA incorporated (purple). Left, raw data converted to ΔpH is shown normalized based on the average of the baseline over the first two minutes (n=3 independent trials). Right, rates obtained by fitting the first 120 seconds after valinomycin was added (black triangle, left) are plotted as a bar graph. An unpaired t-test between empty liposomes and LQLLx610-proteinA show significant change in proton conduction when the channel is present (p<0.0001). **(f)** Native mass spectrometry of LQLLx610-proteinA refolded in C8E4 detergent. Subunit stoichiometry assigned: red, charge state; purple circles, oligomer state. The native MS spectrum confirmed the molecular mass of the pentameric assembly (67,440 Da) and its monomeric protein subunit (13,480 Da). Monomer helical topology (purple cartoon) in micelle and pentameric assembly (top view, colored ribbons) depicted.

To examine whether the changes to folding and stability are sequence-specific, we tested a variant where the accessory domain sequence is scrambled. The resulting LQLLx610scrambled proteins as SUMO-fused (21 kDa) and minimal TM (7 kDa) versions both elute significantly later in SEC than the respective designed pentamer versions (**Fig. S6e,f**), indicating formation of higher molecular weight aggregates. Introducing the random TM span instead promotes misfolding rather than correcting it. Thus, the *de novo* accessory domain’s designed interactions are needed to achieve its stability enhancement.

### Functional multi-pass assemblies derived from LQLLx610

We next tested if the synthetic 10 TM span assembly can be a scaffold for bioengineering applications via structural expansion in the water-soluble region, focused on organized display of immune epitopes. To do so, a variant of the protein-A (protA) 3-helix bundle domain hosting a grafted helical epitope^37^ (GCN4) was designed to extend from LQLLx610’s c-terminus by a rigid continuous helix linker (**Fig. 3c**). The resulting protein, LQLLx610-protA, is expected to organize the folded immunogen with C5-symmetry at membrane surfaces in a pre-determined spacing.

Upon purification, LQLLx610-protA is observed to stably fold to a single oligomeric state (∼70 kDa expected), spontaneously refolds in detergent from organic solvent, and migrates predominantly in gel electrophoresis as an SDS-resistant oligomer faster than expected (∼50 kDa apparent) (**Fig. 3d, S6g**). To assess whether LQLLx610-protA properly displays the foreign membrane-proximal epitope as expected, we used SEC and cryo-EM. The corresponding antibody fragment binds, given its co-elution with oligomeric LQLLx610-protA, confirming conformationally correct display of the helical epitope (**Fig. S7a**). 2D classes of the purified membrane protein-scFv complex further support this, based on observation of multiple scFvs above the micelle in likely pentameric configurations (**Fig. S7b**). Thus, LQLLx610 provides an amenable scaffold for fold expansion. This highlights the potential of engineering membrane proteins customize spacing and multiplicity of domains at membrane surfaces, including to manipulate immune presentation and clustering.

To test whether the LQLLx610-protA assembly retains its function, we tested its ability to conduct protons using a vesicle flux assay. In the absence of an electrochemical driving force, LQLLx610-protA maintained pre-established sodium and potassium gradients (**Fig. 3e**), but Valinomycin-induced potassium efflux, generated an electrochemical gradient that drove proton flux through LQLLx610-protA (**Fig. 3e**) demonstrating high selectivity for protons. After accounting for proteoliposome incorporation efficiency, conduction rates were comparable to the 1-pass LQLL peptide (**Fig. S7c**).

Finally, we tested whether the re-engineered LQLLx610-protA maintains specific pentameric folding, confirming the basis of its proton conductivity. By native mass spectrometry of protein refolded in tetraethylene glycol monooctyl ether (C8E4), the dominant species correspond to pentamer charge states (**Fig. 3f**). These results show that large (70 kDa) functional multi-pass membrane proteins of increasingly complexity can be engineered by sequential synthetic fold expansion, based on minimal *de novo* (3 kDa) TM domain interactions as reliable modular structural building blocks.

### Steric trapping shows the TM accessory domain adds folding stability

To quantify whether the designed TM accessory domain adds stability to LQLL’s pentamerization under non-denaturing conditions, we employed steric trapping^38–42^ in dodecylmaltoside (DDM) micelles (**Fig. 4**). In this assay, solvent-exposed cysteines are biotinylated at sites proximal in the folded complex, such that binding of multiple monovalent streptavidin (mSA, 52 kDa) is sterically restricted until unfolding. As a result, mSA binding is thermodynamically coupled to protein unfolding. While the first 2–3 mSA bind freely (Δ*G*°_Bind_)^39^, additional binding events at remaining sites are hindered until pentamer dissociation (Δ*G*°_D_), producing a biphasic binding isotherm^38,42^. Attenuation of this second phase reports on stability (Δ*G*°_D +_ Δ*G*°_Bind_). mSa binding is quantified via quenching of pyrene-labeled biotin (BtnPyr) upon sequential association with dabcyl-labeled mSA^38^ (**Fig. 4a**).

**Figure 4.**
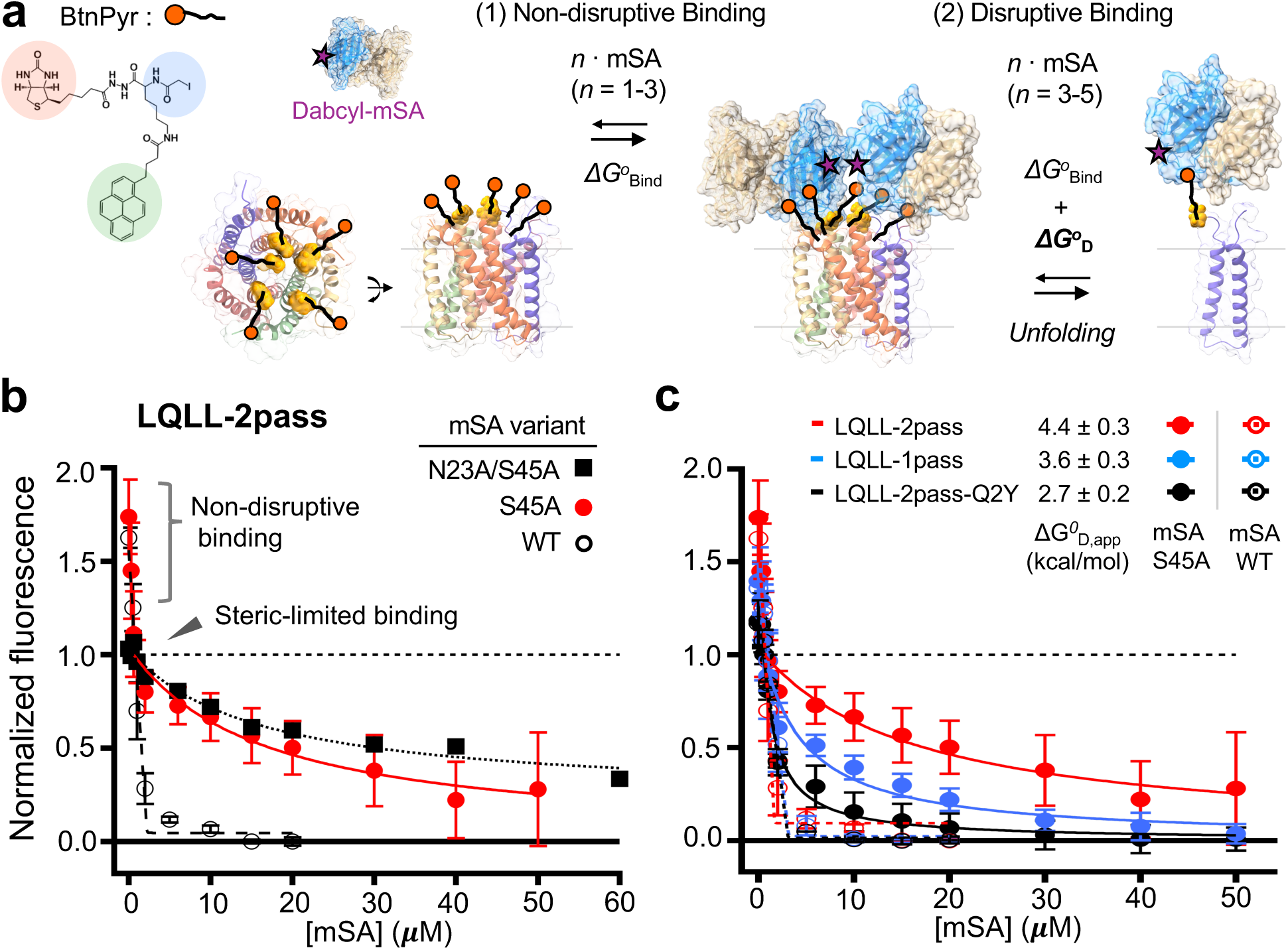
Steric trap unfolding reveals pentameric stabilization by the *de novo* TM accessory subunit. All LQLL constructs used for steric trapping contain an N-terminal SUMO fusion (the monomer molecular weight: 21 kDa). **(a)** A thiol-reactive biotin derivative with fluorescent pyrene (BtnPyr; top-left)^38^ was conjugated to the LQLL-2pass variant with a cysteine at the N-terminus of the pore-lining TM span (P4C, bottom-left). (Right) The reaction scheme for sequential monomeric streptavidin (mSA) binding and steric trapping of the dissociated protein state, used to measure the apparent stability (Δ*G*°_D,app_) of designed LQLL pentamer variants. The first few events of mSA binding to biotin linked to each pentamer are not sterically limited (∼2-3 mSA molecules, Δ*G*°_bind_). Subsequent mSA binding is sterically occluded, thus energetically coupled to dissociation of TM domain pentamer. **(b)** Binding isotherms of LQLL-2pass (fixed at [monomer] = 1.0 μM) in 2 w/v-% DDM micelles at room temperature with mSA variants of different intrinsic biotin affinities: *K*_d,biotin_ = 2ξ10^−7^ M (mSA-N23A/S45A), 9ξ10^−9^ M (mSA-S45A), and ∼10^−14^ M (mSA-WT)^38,42^. Binding was monitored by quenching of BtnPyr fluorescence on LQLL-2pass by dabcyl-conjugated mSA. In each isotherm, fluorescence intensities (dots) were normalized to the amplitude of the attenuated second binding phase. Errors denote SEM (*N* = 3 biological replicates). Weakened mSA variants display a rapid first phase of binding events followed by an attenuated second binding phase ([mSA] > ∼0.6 μM), reflecting mSA binding that is limited by pentamer association-dissociation events. **(c)** Binding isotherms with mSA-S45A comparing pentamer variants LQLL-2pass (red), LQLL-1pass (blue), and control clash-inducing mutant with impaired pentamerization LQLL-Q2Y-2pass (black). Dots represent pyrene fluorescence intensities; lines represent best-fit binding curves. The fitted Δ*G*°_D,app_ values are listed. Dashed lines with empty dots are binding isotherms with mSA-WT displaying maximal fluorescence quenching (i.e., saturated binding) levels. Errors denote ± SEM (*N* = 3 biological replicates).

After screening (**Fig. S8**), we identified a suitable biotinylation site at the N-terminus of LQLLx610-2pass’s inner TM helix where upon mSA binding titration led to a saturated and biphasic isotherm (**Fig. 4b**). The apparently sterically hindered second binding phase began at approximately 0.4-0.6 molar equivalents (∼2-3 mSA) as expected. To achieve suitably attenuated binding to biotinylated LQLLx610-2pass, a higher-affinity variant of mSA (S45A, *K*_d,biotin_ = 9ξ10^−9^ M) was required compared to the variant needed to capture glycophorin-A dimer unfolding (mSA-S27R, K_d,biotin_ = 6.5×10^−6^ M), confirming that pentameric folding is strong^38,42^. SDS-PAGE mobility-shift^39^ confirmed that binding of mSA indeed induces pentamer dissociation (**Fig. S8d**).

Next, we compared the relative stabilities of the designed LQLLx610-2pass with the original LQLL-1pass and a variant LYLLx610-2pass hosting a pentamer-disrupting “Q2Y” core mutation (Q14ΔY) as a negative control (**Fig. 4c, S9**). Amongst these, LQLLx610-2pass displayed the strongest steric hindrance to unfolding given the extent of attenuation in the second mSA-binding phase. Because no analytical solution is available for the mSA-induced pentamer–monomer equilibrium and since intermediate oligomeric states may be populated, we could not reliably determine a true free energy of pentamer dissociation. Instead, to compare these variants, we modelled the attenuated binding phase as monomeric protein denaturation^38,41^ to derive relative apparent free energy changes for dissociation (Δ*G*°_D,app_). The designed LQLLx610-2pass, Δ*G*°_D,app_ = 4.4 ± 0.3 kcal/mol, is significantly stronger than the LQLL-1pass’s intermediate stability (Δ*G*°_D,app_ = 3.6 ± 0.2 kcal/mol). The LYLL-2pass-Q2Y pore mutation substantially weakens pentamerization (Δ*G*°_D,app_ = 2.7 ± 0.2 kcal/mol) further supported by the loss of SDS-resistant pentamer on SDS-PAGE. However, this mutant is not fully monomeric given observed steric hindrance (**Fig. 4C, S9a**). These results corroborate that our design strategy to incorporate synthetic outer-shell TM helix indeed significantly bolsters folding under non-denaturing conditions as intended.

### *De novo* fold expansion to large single-chain multi-pass proteins

Although minimal self-assembling peptides serve as foundational units in evolution, longer continuous chains are better suited to explore functional sequence space – able to asymmetrically modify structure and function, such as within channel pores. This notion motivated us testing whether a folded single-chain 10-span membrane protein could be engineered from the *de novo* pentameric 2-pass subunits, analogous to natural gene fusion (**Fig. 1b, 5a**). The 1-pass LQLL pentamer of parallel helices is not amenable to direct single-chain fusion due to distant inter-subunit termini. Instead, incorporating the outer-shell antiparallel helices of the accessory TM subunits provide tractable linkage points. Design of four helix-turn-helix junctions (5-13 residues, largely solvent-exposed) joining accessory domain C-termini to the channel-lining N-termini allows sequential connections between subunits. While the pentamer of 2-pass LQLLx610 subunits is not predicted by AF3 (**Fig. S10a**), versions of the resulting 10-pass single-chain “LQLLx610pass” protein are confidently predicted within 1.2 Å RMSD to the design model (**Fig. 5b, Table S4**), indicating this structure is most likely the lowest energy fold for the sequence.

**Figure 5.**
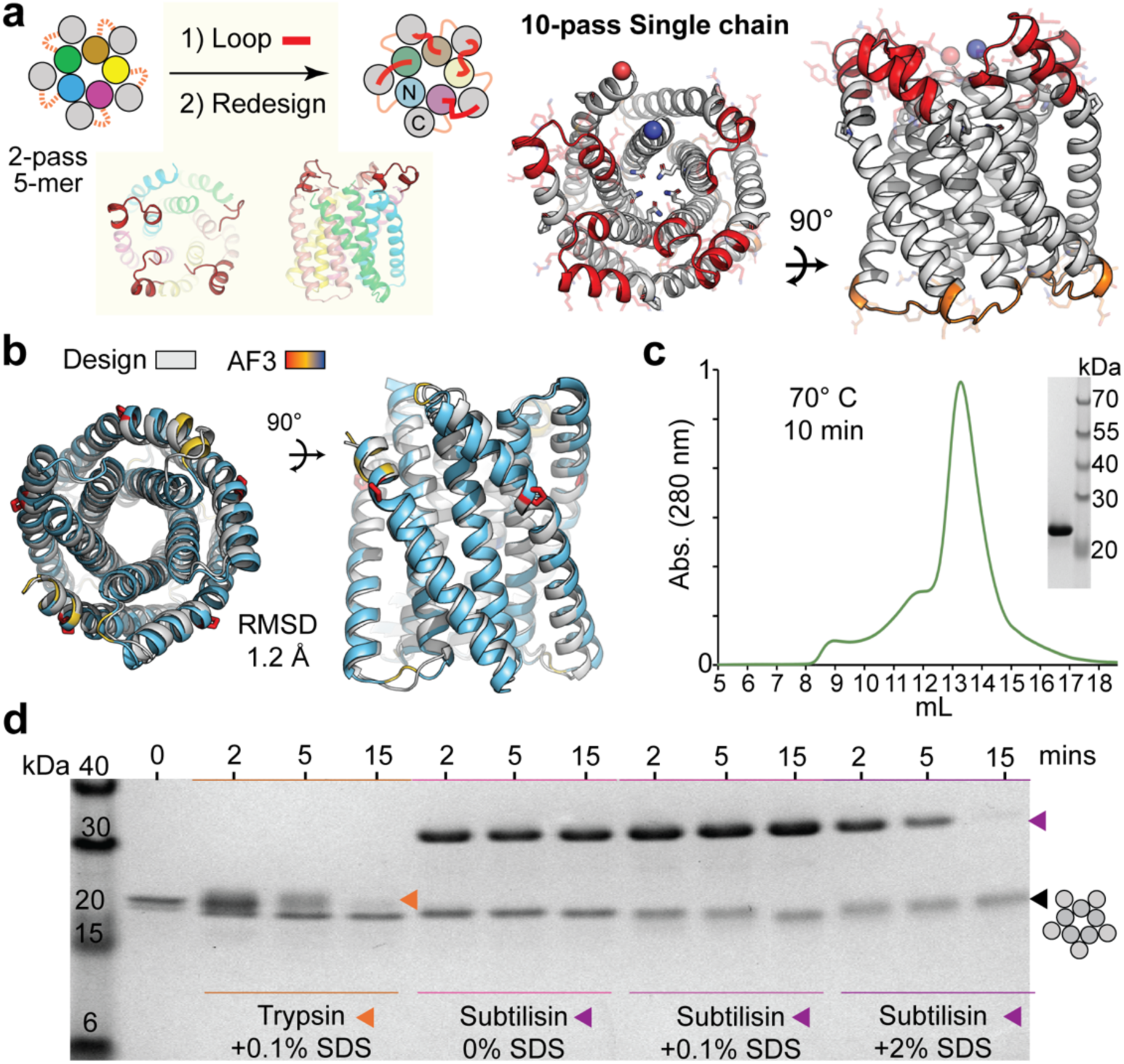
Fold expansion and stability of *de novo* 10-pass channel variant LQLLx610pass. **(a)** Left, modeling new loops (red) connecting the 2-pass pentamer (5-mer) *de novo* TM subunits and redesign of those remodeled regions, resulting in monomeric single-chain 35 kDa LQLLx610pass. Right, side and top views; C-and N-termini as red or blue spheres, respectively; pore glutamine ring sidechains as sticks. Orange, loops retained from 2-pass pentamer LQLLx610. **(b)** AF3 predicted structure (colored by pLDDT) superposed with LQLLx610pass* design model (gray). Designed proline kink in *de novo* accessory TM span, red sticks. pTM: 0.84 **(c)** SEC UV chromatogram of LQLLx610pass in 1 mM DDM mobile phase after 30 minute incubation at 70 °C, showing resistance to thermal-induced aggregation, representative of n=3 trials. Inset, SDS-PAGE **(d)** Resistance of LQLLx610pass to proteolysis at 55 °C, read by SDS-PAGE. Protein at 0.1 mg/ml in 5 mM DDM incubated at 55 °C with 0.2 mg/mL trypsin or subtilisin enzyme (protease bands, orange or purple markers, respectively) incubated with 0, 0.1, or 2% w/v SDS.

Upon purification, LQLLx610pass was highly stable and ran on SEC as a single monodisperse species as the same size as the 2-pass pentamer in all detergents tested (**Fig. 5c**). Its migration on SDS-PAGE matches that of pentameric versions (observed: ∼25 kDa; expected: 35 kDa). In addition to resistance to heat-induced aggregation (70°C, **Fig. 5c**), LQLLx610pass is highly resistant to proteolytic degradation (trypsin, subtilisin) under harsh conditions known to unfold and rapidly digest bacteriorhodopsin^43^ (55° C, with SDS) (**Fig. 5d**). It is important to note that the extra-membrane polar regions bridging TM spans are long enough that, if unfolded, would offer protease-accessible cleavage sites. Significant degradation is only observed with high SDS and prolonged exposure (2%, 15 minutes) yet remains incomplete. These behaviors suggest the monomeric single-chain protein adopts a highly stable compact folded structure as designed.

Next, we assessed the 10-spanning TM proteins for trafficking in mammalian cells to test their utility as scaffolds in cellular engineering applications. To encode an topology of an extracellular N-terminus (“N-out”), an expression construct with a cleavable ssHA signal sequence preceding the minimal 2-pass pentamer was transfected to HEK293T cells. Unpermeabilized cell immunostaining reveals abundant synthetic membrane protein expression with correct trafficking and N-out topology at the plasma membrane surface (**Fig. S11a**). Then, we further tested the single-chain LQLLx610pass for its malleability to altering insertion topology as an mScarlet fusion. We directed the N-terminus either intracellularly (“N-in”) using a previously identified non-cleavable polar ssPolar signal sequence^35^ (**Fig. S11b**) or extracellularly (“N-out”) using a hydrophobic ssHA (**Fig. S11c**). Fluorescence microscopy revealed distinct trafficking patterns. The “N-out” construct showed higher expression and partial plasma membrane localization but was mostly found in intracellular puncta, whereas the “N-in” construct was more evenly distributed, outlining cell edges, indicating stronger membrane localization. These results show that trafficking of these *de novo* multi-pass proteins can be manipulated within human cells, and highlight their amenability to navigate cellular machinery and complex mammalian membranes as engineering-compatible scaffolds.

### Cryo-EM structure of LQLLx610pass confirms intended design

To test whether the *de novo* designed TM accessory subunit interacts within the inter-subunit groove and stabilizes the pentameric core as anticipating, we generated a LQLLx610pass construct with a fiduciary marker for structure determination^44^. A synthetic water-soluble helical bundle, Lockr (35kDa, pdb: 7JH5^45^), was incorporated between TM helices 8 and 9 through design of two continuous rigid linker helices. The resulting LQLLx610pass-Lockr89 protein was confidently predicted by AF3 to adopt the intended fold (**Fig. S10c**) and is expressed in mammalian cells (**Fig. S11d**). To further increase particle size, helical epitopes were installed at two solvent-exposed sites on Lockr, which successfully facilitated stable complexes with scFv fragment antibodies^37^ (**Fig. S12a-b**). LQLLx610pass-Lockr89 is stable, monomeric, and well-behaved when purified in DDM and glyco-diosgenin (GDN) (**Fig. S12a**). The cryo-EM structure of it 2:1 scFv:protein complex was solved, having local resolutions of ∼3 Å in the soluble domain and ∼5 Å amongst TM spans (**Figs. S12c-d, S13**).

The protein’s overall fold is unambiguously discerned from the map (**Fig. 6a**) and is in strong agreement with the expected 10-pass pseudo-symmetric channel fold, with the structural model fitting well into the density. Densities for 4 of 5 designed accessory helices (TMs: 2,4,6,8) are accurately positioned within the inter-subunit groove adopting the helical packing geometry intended, with TM10 too poorly resolved. ScFv molecules were observed engaging the helical epitopes site grafted onto Lockr as expected (**Fig. S14a-b**), validating the fiduciary marker strategy. The density shows strong bending in accessory helices, best resolved in TM8 by a ∼33° kink facilitate its packing to pore-lining TM7 and TM9 at the site where Pro263 is located within the model to as designed (**Fig. 6b**). Thus, the experimental structure supports that both the correct fold expansion and intended mechanism of stabilization driven by the accessory TM domain were achieved, including designed helical distortions.

**Figure 6.**
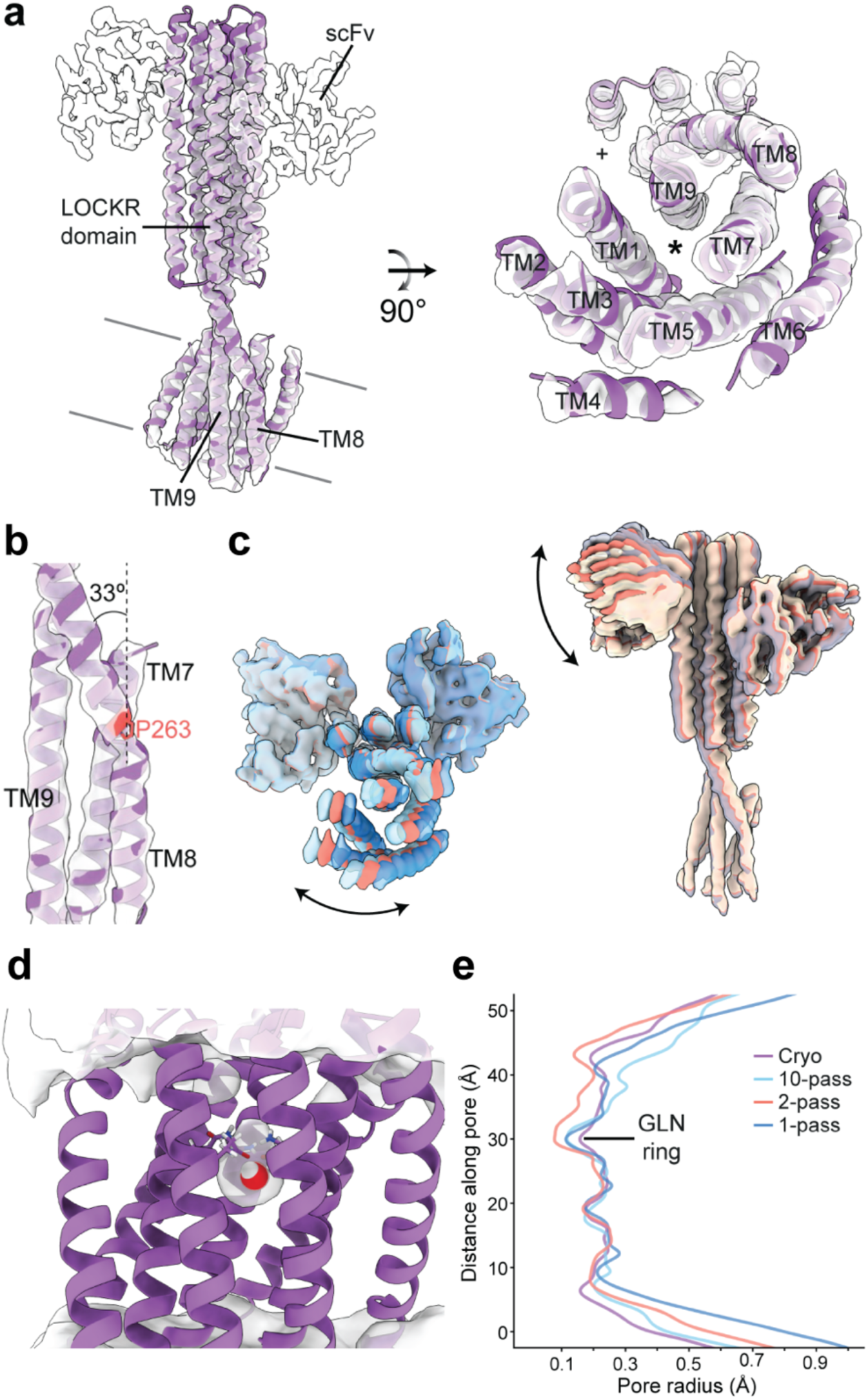
Cryo-EM structure of LQLLx610pass Lockr fusion. (**a**) Left, cryo-EM density for LQLLx610pass-Lockr fusion and fitted model. Right, close-up view of the TMD, showing density for 9 out of the 10 designed TM helices. (+) depicts expected localization of TM10. (*) depicts expected position of the pore pathway. **(b)** Kink in TM8 induced nearby P263 (red), allowing TM8 to contact and stabilize both TM7 and TM9. **(c)** Densities obtained from 3DFlex analysis along two dimensions in latent space. Left, densities for first dimension depict coordinated motion of the TMD helices inside the micelle. Right, densities for second dimension show tilting motion of Lockr relative to the fixed micelle, most visible in one scFv. **(d)** Frame of an MD simulation for cryo-EM structure of LQLLx610pass-Lockr illustrating localization of water molecules around the pore pathway. Density represents average position of water molecules across a 1μs simulation. Glutamine ring selectivity filter shown as sticks. One water molecule present in this frame is shown as spheres. TM4 is removed for visibility. **(e)** Comparison of pore profiles for different LQLL designs: LQLLx610pass-LOCKR89 model fit into cryo-EM structure density (purple), LQLLx610pass design model (light blue), LQLLx610-2pass pentamer design model (red), and LQLL 1-pass pentamer X-ray structure (dark blue, PDB:7UDZ).

Given the sizable difference in resolution between the Lockr and LQLLx610pass domains, we performed 3DFlex analysis to discern dynamics within structure (**Fig. S12e**). Two main motions were identified (**Fig. 6c**): rotation of the channel TM helices within the micelle relative to Lockr and tilting of Lockr with scFvs relative to the TM bundle and micelle (given the imposed rigid micelle in 3DFlex). Together, these results indicate Lockr and LQLLx610pass move significantly as independent domains relative to each other, indicating the extended 2-helix linker region is more flexible than expected and likely limits TM helix resolution.

While the folds are consistent overall, we analyzed pore characteristics to assess whether channel structures were altered in multi-pass LQLLx610 variants. Amongst single and multi-pass LQLL channels, pore radial profiles are nearly identical within the apolar section, with deviations observed mostly at pore vestibules (**Fig. 6c**). Although the glutamine (Q)-ring selectivity filter appears slightly more open in LQLLx610pass-Lockr89, this cannot be confidently concluded at the current resolution. MD simulations of this experimental structure result in pore hydration and constriction similar to that previously observed^18^, including strong occupancy of a single water at the Q-ring (**Fig. 6b**). Thus, multi-pass variants maintain similar channel chemistry at the selectivity filter, expected to stabilize transient water wires and selective proton transport.

### Conclusions

While design of helical multi-pass structures has progressed tremendously^10–13^, most design strategies have relied on either water-soluble proteins as templates, known TM sequence patterns^28,46–48^ (e.g. Small-X6-Small)^34^, or extra-membrane globular domains to stabilize *de novo* TM structures^10–13^. Here, we demonstrate a strategy focused on membrane-specific interaction principles and design of modular building blocks sequentially to reliably construct *de novo* multi-pass proteins of increasing size and complexity. Adding “accessory-stabilizer” outer-shell TM helices to a “target” inner-shell core result in topologies resembling natural pentameric ligand-gated^49^ and synthetic voltage-gated channels^12^: with outer helices buttressing pore-forming TM span conformations. Our approach utilizes a strategy similar to the “rationally-seeded” design approach, which expanded peptide coiled-coils into large single-chain water-soluble helical bundles^50^, but diverges in implementation and design principles to achieve this in membranes. These *in silico* template-based fold expansion methods mirror how natural proteins can evolve from smaller modules via sequential domain fusion and diversification^51,52^. Likewise, the accessory TM domain design strategy is immediately relevant for membrane protein engineering. Its application as “folding corrector” fusion domains could bolster the intended structures sought in many recent design attempts of single-spanning helical nanopores or synthetic receptors where folding was found to be marginally stable, nonspecific, or adopting unintended architectures^28,46,47,53,54^.

Our work, alongside that of recently reported 2-pass GPCR-modulators^15^, marks advances in *de novo* design of interaction with folded multi-helical TM conformations, moving beyond the previous cutting-edge limited to complexes of simple single isolated TM helices^22,23^. To this end, we implemented an emerging vdW packing criteria^34^ for apolar steric specificity alongside data-driven design approaches to mine binding template structures and interaction principles directly from observed membrane protein structures. Notably, the target-conditioned method converged on designs utilizing kinked helices, a molecular feature common to natural TM spans^55^ but not yet explicitly incorporated or noted as a key feature in membrane protein design to date to our knowledge. Interestingly, directed evolution efforts have resulted in improvement of protein catalytic functions through inclusion of a proline-induced kink^56^, indicating the engineering value of precisely positioning helices. Our design proactively navigated the challenge to stabilize a local helix distortion and achieve specific helix packing angles by incorporating sufficient supportive favorable interactions like the key expected sidechain H-bonds tailored to the kinked geometry (**Fig. 2b**, **S2b**). High-confidence AF3 predictions and the electron density suggest successful incorporation of this feature by these design principles (**Fig. 6a-b, S3, S14**).

The lead TM domains demonstrated capacity to bind the pentamer complex *in vitro* as separate peptides, but only weakly, indicating affinity is the critical area of improvement required to achieve biologically relevant “folding corrector” applications. Nonetheless, upon fusion, the *de novo* accessory TM domain interactions were confirmed to provide impactful energetic stabilization (∼1 kcal/mol, typical for a single TM helix interaction) which significantly improves ion channel folding. While the underlying atomic-level interactions are unclear at the electron density’s resolution, the mechanism of TM stabilization is consistent with our design model within the intended binding groove. Despite successfully encoding the kinked structure, inclusion of proline and distorted helices within our design may have been to the detriment of the overall interaction strength. Expanding the protein mass and interaction interface by design of additional TM or soluble domains may be other strategies to improve affinity. However, our minimal single-helix strategy is advantageous given potential to deliver as synthetic peptides to cells and tissue, offering a route to “folding corrector” applications without requiring genetic delivery of larger multi-pass constructs (e.g., mRNA or AAV). Here, our proof-of-concept validates the success of this design method for generating interactions to straight ideal α-helices of a model ion channel. For computational strategies to be generalizable for creating new pharmacological chaperones stabilizing disease-relevant human proteins, they should be apt to encode interactions with the kinked irregular dynamic TM helix conformations prevalent throughout the membrane proteome. As a complement to emerging diffusion-based approaches, our work here demonstrates tools and design guidelines that serve as vital steppingstones to unlock polypeptide-based strategies to combat pathological misfolding of membrane proteins on-demand.

## Acknowledgements

A portion of this research was supported by NIH grant R24GM154185 and performed at the Pacific Northwest Center for Cryo-EM (PNCC) with assistance from Marcelo de Farias and Sean Mulligan. N.P.J. was supported by the Molecular and Cellular Biophysics NIGMS T32 training grant T32GM148376. H.T.K. acknowledges support from the NIH NIGMS R00GM138753. J.R.Y acknowledges funding from the NIH (R01-AG075862-02, R01-HL165168-01, R01AG077046-02). H.H. acknowledges support from NIH R35GM144146. M. Murakoso was supported by the Olson-King Fellowship in Computational Biology. N.F.P acknowledges funding from NIH (R00GM135519) and the Innovation Research Fund of Dana-Farber Cancer Institute to support this work.

---

**Supplementary Figure 1.**
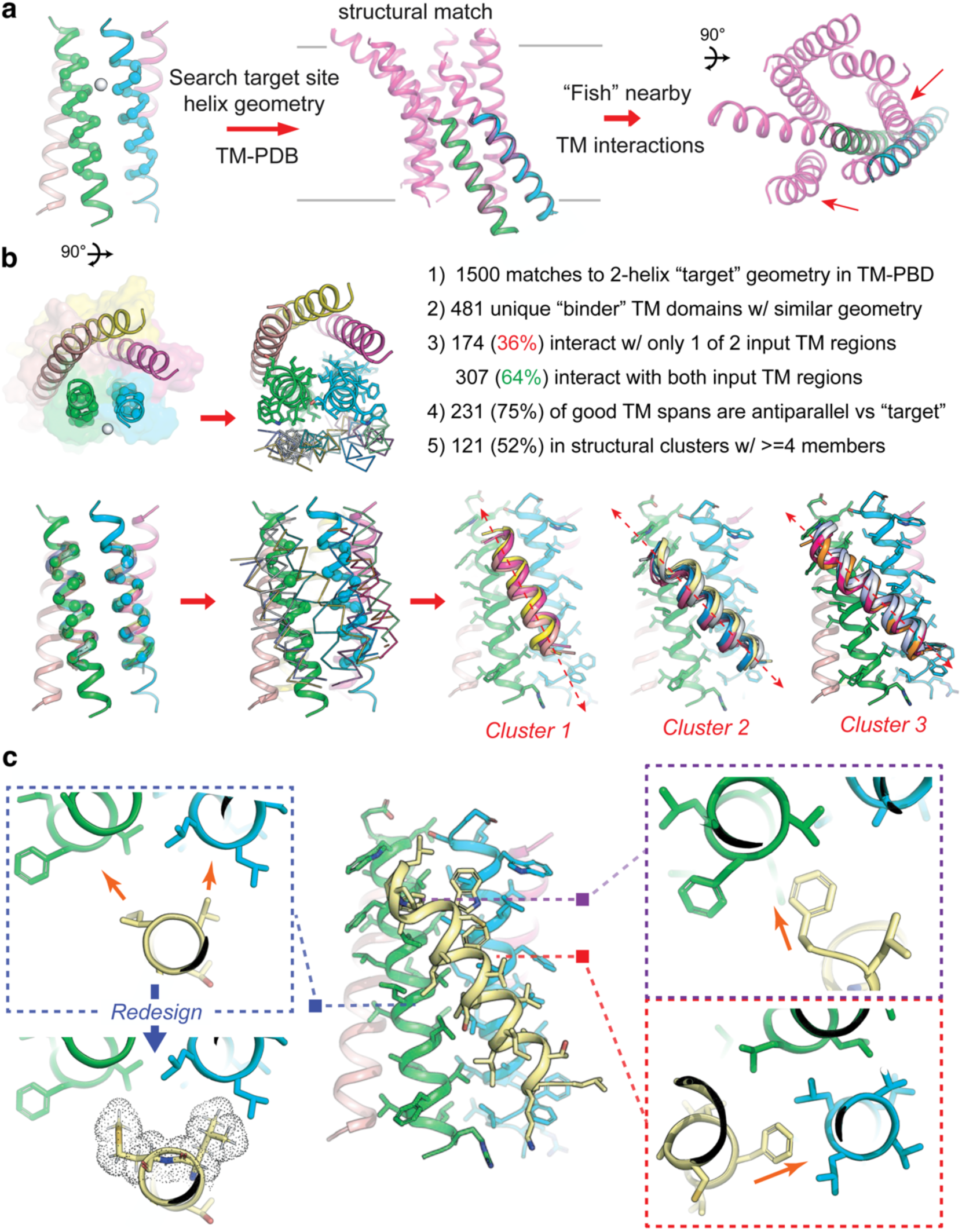
Model building *de novo* adaptor subunit TM helix. (**a**) Left, eVgL pentamer “target” intended binding surface site marker by white sphere; C-alpha atoms (spheres) denote the 15-residue region of 2 adjacent sub-unit helices queried for structurally similar instances in other membrane protein folds. Middle, backbone geometry search (MASTER) example of eVgL TM domain interface (cyan/green helices) matching to sub-structure of a multi-pass membrane protein (pdb: 5EE7; human glucagon receptor, purple). Right, “fishing” new instances of TM domains packing into the input 2-helix geometry (red arrows), finding those backbone tertiary arrangements compatible with interacting with eVgL’s subunit interface. **(b)** Left, overlay of new TM domain backbone geometries (color, ribbon helix trace) fished from membrane structures, aligned to the 2-helix search query. Top right, 481 compatible TM helix geometries found from ∼1500 low RMSD matching instances of the input query. Of those 307 extensively contact both helices. An antiparallel preference is observed (235/307). Bottom right, fished antiparallel TM domains structurally cluster into variations of a consensus interacting geometry – informing our backbone positioning of TM accessory subunit. **(c)** Example of a fished TM domain (yellow) extracted from the sphingosine receptor (TM6, pdb: 3V2Y) and aligned with the targeted eVgL binding region, which exhibits privileged interaction geometry. Sidechain directions (orange arrows) are pre-disposed for favorable direction into helix “holes” for knobs-into-holes packing. Right, purple and red lined inset, the native sidechains of the fished TM interaction without design make compatible interactions with eVgL’s targeted binding surface. Left inset, blue box, sidechain vectors are directed towards helix holes, and protein design can improve packing interactions to stabilize the complex templated by the modelled TM domain interactions.

**Supplementary Figure 2.**
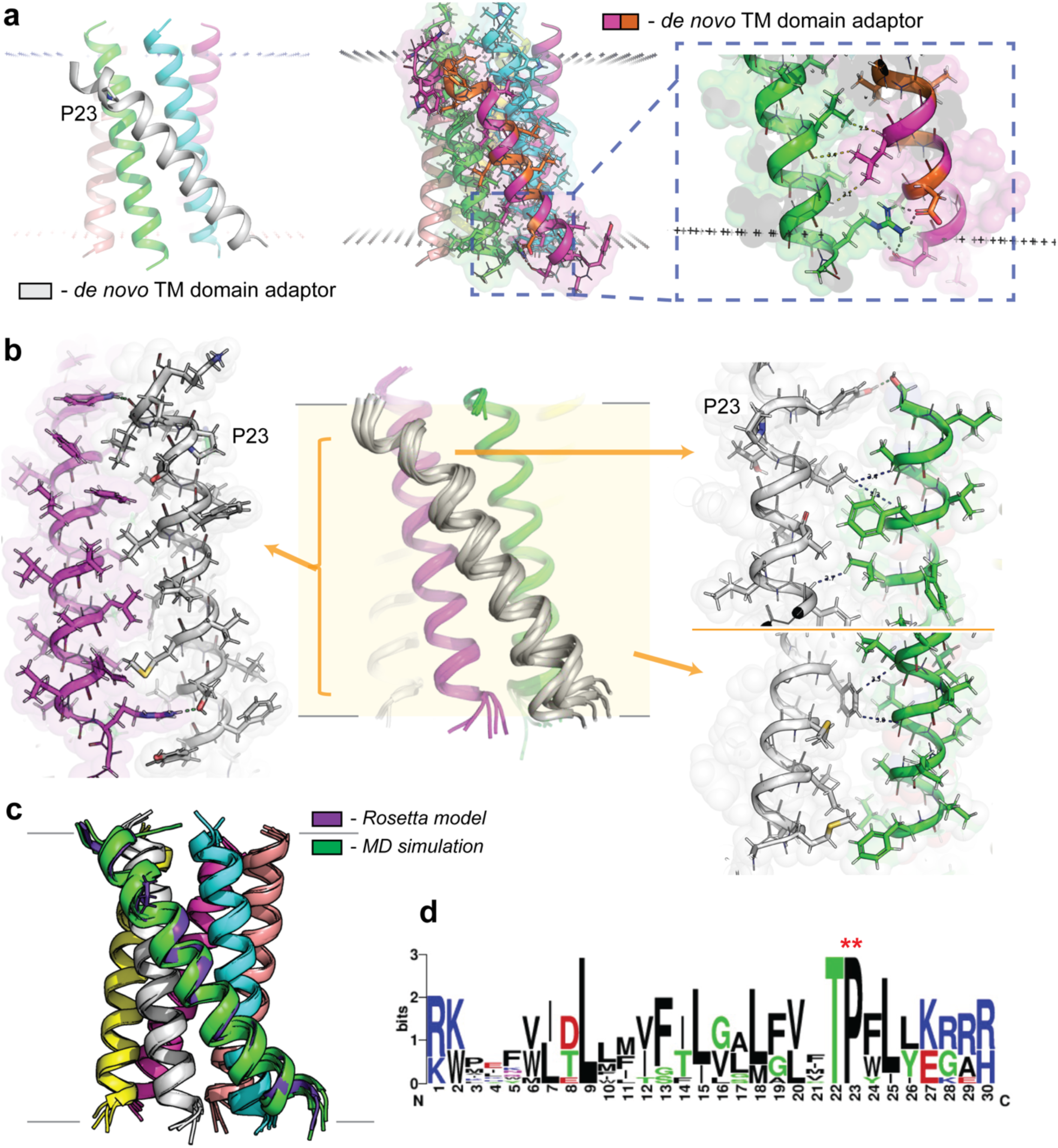
Design of *de novo* TM adaptor subunits. (**a**) Left, modelled TM accessory subunit (white) in interaction complex with eVgL, noting consensus proline conserved from the clustered natural interacting TM domains. Right, example all-atom relaxed design model (orange and purple *de novo* TM adaptor. Blue inset, shows key backbone-directed apolar sidechain packing with close contacts (<3.5 Å) and salt bridge network at the lipid-water headgroup interface region of the membrane. Black “+” markers denote implicit membrane. **(b)** Center, overlay of backbone (white cartoon) of *de novo* TM subunit MD simulation frames of design TM483 aligned to eVgL pentamer structure, including proline kink. Left, extended view of apolar packing interface and hydrogen bond (replacing salt bridge in panel a) with eVgL TM subunit 1 (purple). Right, interactions of close apolar sidechain backbone contacts with eVgL TM subunit 2 (green). **(c)** Backbone overlay of interaction complex from design (purple helix cartoon) compared to MD simulation frames (green) for design TM610. **(d)** Sequence logo of de novo TM adaptor subunits final design models (Table S1), showing strong diversity in interactions. Conserved proline and i-1 threonine motif stabilizing the helix kink where fixed throughout design.

**Supplementary Figure 3.**
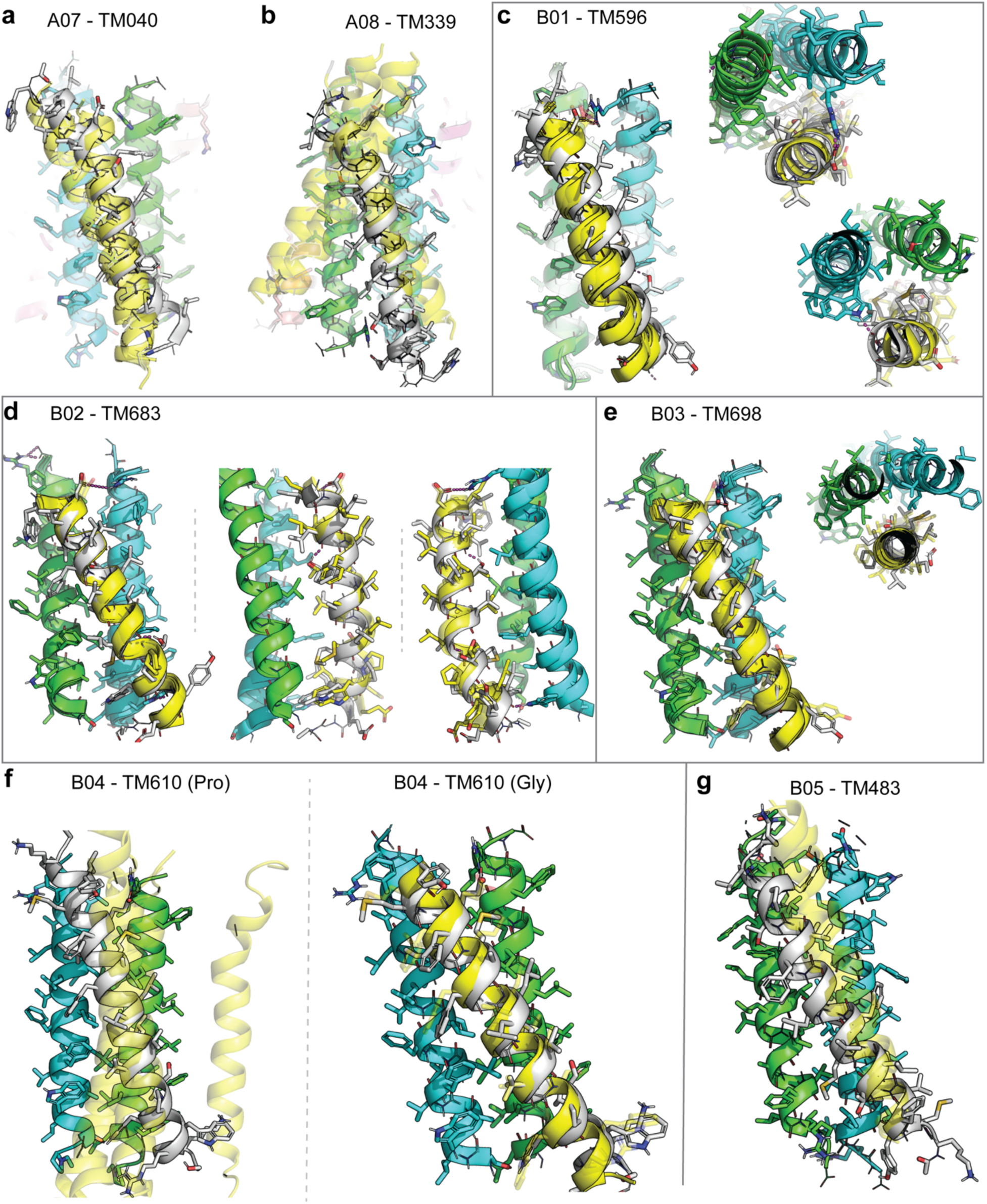
AlphaFold prediction of TM adaptor binding to target interface of pentamer eVgL interface. (**a**) Overlay of TM adaptor helices from the intended design model A07-TM040 (grey, cartoon; sidechains as sticks) versus AF3 predicted conformations (transparent yellow, top 3 models), relative to the targeted binding site on the surface of eVgL comprised of helices (blue, green). **(b)** Design model of A08-TM339 (gray) versus AF3 prediction (transparent yellow, top 5 models). **(c)** Left, design model of B01-TM683 (gray) versus AF3 prediction (yellow). Middle, top down of c-terminus the pentamer. Right, top down of n-terminus the pentamer. Interhelical hydrogen bonds denoted as pink dashes. **(d)** Left, design model of B02-TM339 (gray) versus AF3 prediction (transparent yellow, top 5 models). Middle, side view of protomer A interface (green). Right, side view of protomer B interface (blue). **(e)** Design model of B03-TM698 (gray) versus AF3 prediction (transparent yellow, top 3 models). Right, top down view. **(f)** Design model of B04-TM610 (gray) versus AF3 predictions (transparent yellow, top 3 models). Left, proline variant of TM610; right, glycine variant of TM610. **(g)** Design model of B05-TM043 (gray) versus AF3 prediction (transparent yellow, top 3 models).

**Supplementary Figure 4.**
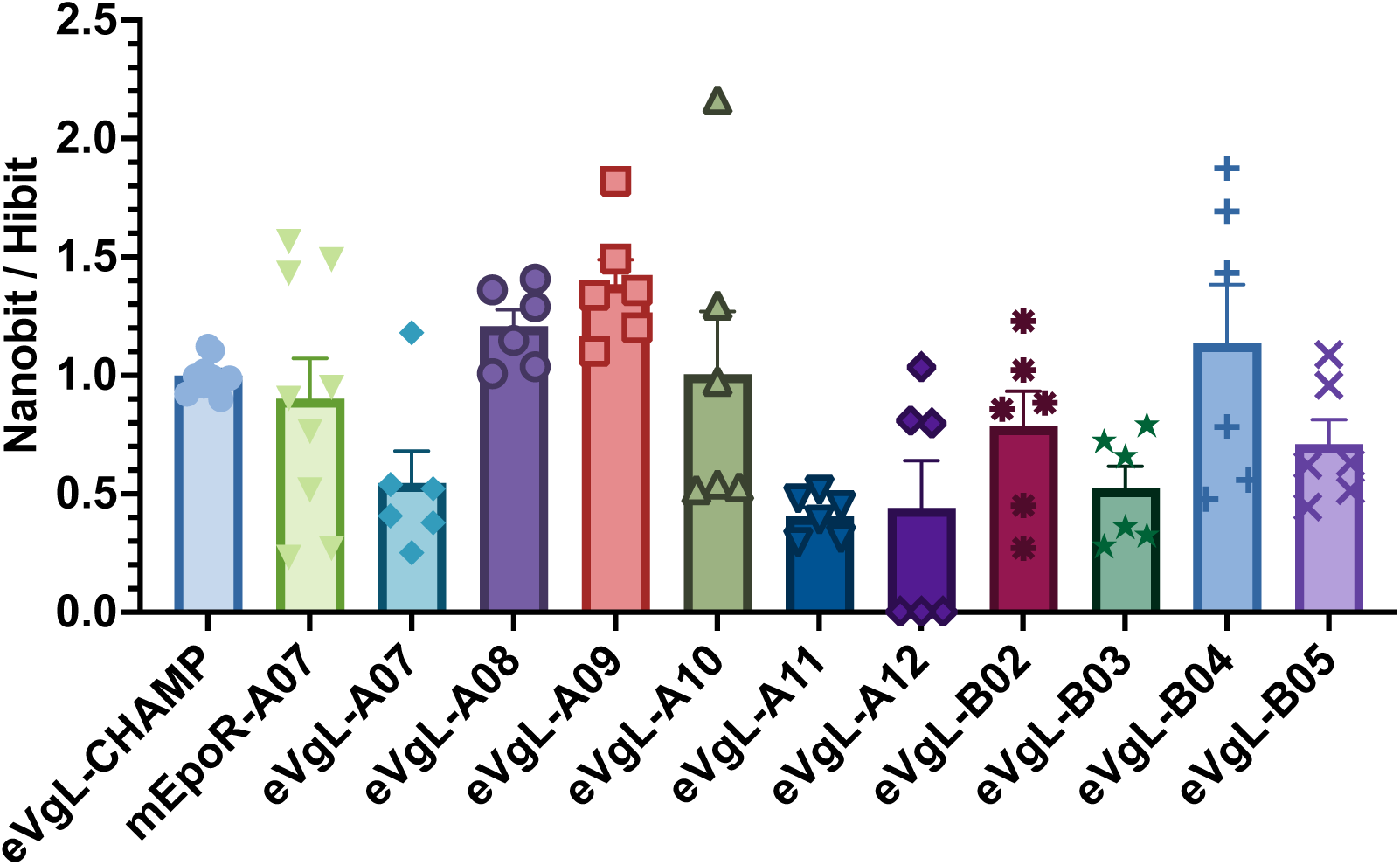
NanoBit membrane protein-protein interaction (PPI) measurements. NanoBit luminescence measurements of PPI interaction upon co-expressing pairs of LgBit “binder” proteins and SmBit “target” proteins by transient transfections in HEK293T cells. The interaction propensity is given (y-axis) after normalization of the NanoBit PPI signal by a second luminesce measure of the total expression of the binder (as determined by addition of HiBit and reading the LgBit-HiBit reconstituted enzyme signal). eVgL-CHAMP and mEpoR-A07 are negative controls. Other pairings are binders design against eVgL (Table S3). Scatter points represent 2-3 biologically independent trials with n = 3 technical replicates. Each biological replicate dataset was normalized to the average of eVgL-CHAMP control group of the same biological replicate.

**Supplementary Figure 5.**
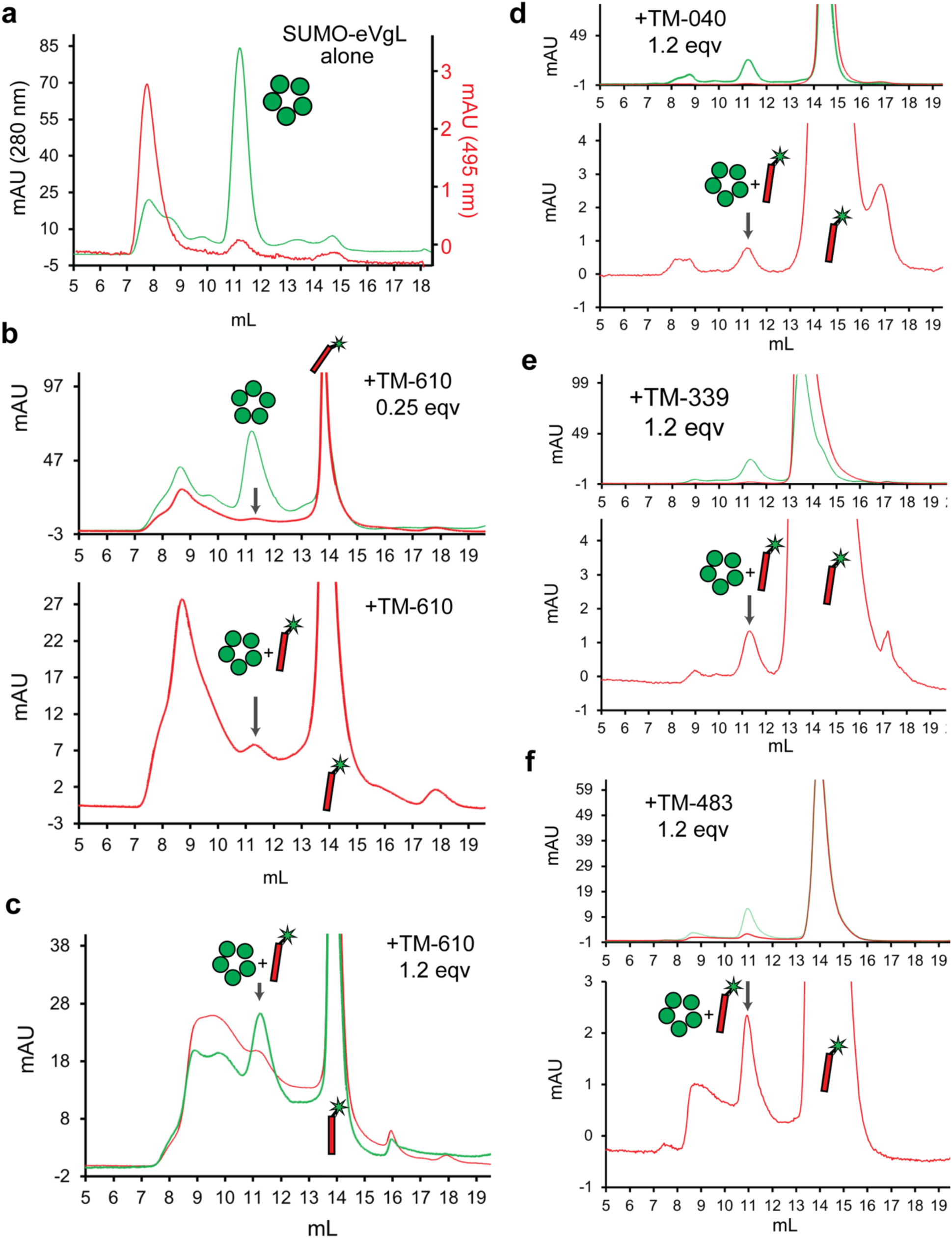
Size exclusion chromatography of fluorescein-labeled synthetic TM subunit peptides with pentameric SUMO-eVgL. (**a**) SEC chromatogram of 280 nm (green trace, left axis) and 495 nm (red trace, right axis) of 2 mg/ml SUMO-eVgL-1pass (green circles, pentamer) in s200i column with 3 mM C14B mobile phase. **(b)** Top, SEC chromatogram overlay of 280 nm and 495 nm traces for 25 μM fluorescein-labelled *de novo* TM-610 (red helix cartoon) and 110 μM (2 mg/mL) SUMO-eVgL-1pass. Bottom, zoom in 495 nm trace, arrow on co-eluting species. **(c)** Top, SEC chromatogram overlay of 280 nm and 495 nm traces for 65 μM fluorescein-labelled *de novo* TM-610 and 50 μM (1 mg/mL) SUMO-eVgL-1pass. Arrow on co-eluting species. **(d)** Top, SEC chromatogram overlay of 280 nm and 495 nm traces for 65 μM fluorescein-labelled *de novo* TM-040 and 110 μM (2 mg/mL) SUMO-eVgL-1pass. Bottom, zoom in 495 nm trace, arrow on co-eluting species. **(e)** Top, SEC chromatogram overlay of 280 nm and 495 nm traces for 65 μM fluorescein-labelled *de novo* TM-339 and 110 μM (2 mg/mL) SUMO-eVgL-1pass. Bottom, zoom in 495 nm trace, arrow on co-eluting species. **(f)** Top, SEC chromatogram overlay of 280 nm and 495 nm traces for 65 μM fluorescein-labelled *de novo* TM-483 and 110 μM (2 mg/mL) SUMO-eVgL-1pass. Bottom, zoom in 495 nm trace, arrow on co-eluting species.

**Supplementary Figure 6.**
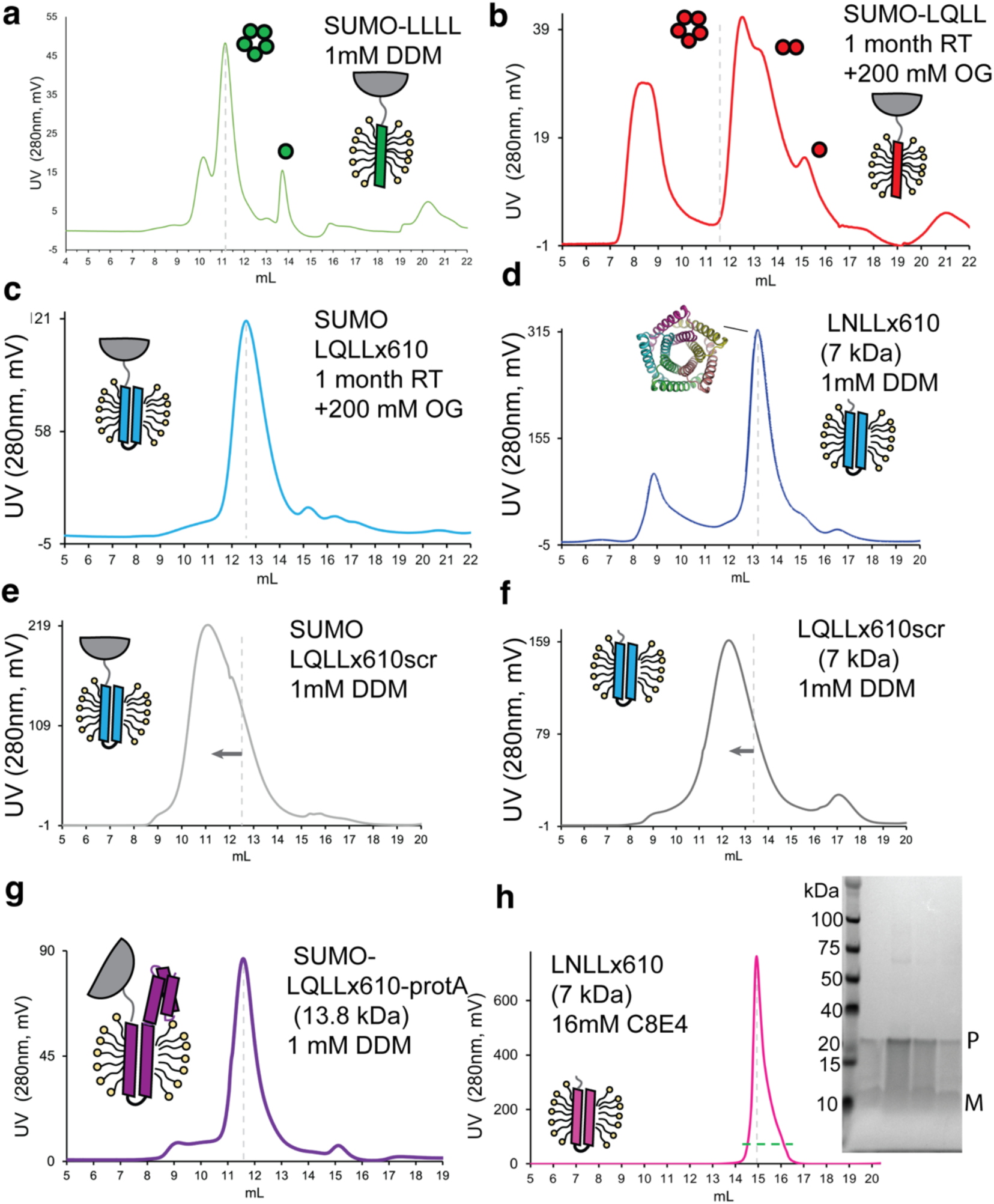
Size exclusion chromatography of SUMO and cleaved pentameric TM domain protein single– and multi-pass oligomeric constructs. (**a**) SEC chromatogram 280 nm of SUMO 1-pass eVgL-LLLL at 2 mg/mL purified from Ni-NTA in C14B injected to 3 mM C14B mobile phase. Representative of n=3 trials; replicate of Figure 2a. Cartoon, topology of monomer. Gray dash indicated pentamer elution. **(b)** SEC chromatogram of SUMO 1-pass LQLL at 2 mg/mL purified from Ni-NTA in C14B, then incubated in 200 mM OG for 1 month at room temperature, then injected to 3 mM C14B mobile phase. Representative of n=3 trials; replicate of Figure 2c. Cartoon, topology of monomer. Gray dash indicated eVgL’s pentamer elution time, showing slight shift for LQLL. **(c)** SEC chromatogram of SUMO 2-pass LQLLx610 at 2 mg/mL purified from Ni-NTA in C14B, then incubated in 200 mM OG for 1 month at room temperature, then injected to 3 mM C14B mobile phase. Representative of n=3 trials. Cartoon, topology of monomer. **(d)** SEC chromatogram of cleaved 2-pass LNLLx610 variant at 2 mg/ml into 1 mM DDM mobile phase. Cartoon, topology of monomer. Representative of n=3 trials **(e)** SEC chromatogram of SUMO 2-pass LQLLx610scr (scrambled 2^nd^ *de novo* TM domain) at 1 mg/mL into 1 mM DDM mobile phase. Representative of n=3 trials. Dashed line denotes elution time of unscrambled SUMO-LQLLx610 elution time, indicating a larger assembly is formed by the scrambled protein **(f)** SEC chromatogram of cleaved 2-pass LQLLx610scr (scrambled 2^nd^ *de novo* TM domain) at 1 mg/mL into 1 mM DDM mobile phase, Cartoon, topology of monomer. Representative of n=3 trials. Dashed line denotes elution time of 2-pass LQLLx610 in panel (d). **(g)** SEC chromatogram of SUMO 2-pass LQLLx610-proteinA fusion at 1 mg/mL into 1 mM DDM mobile phase, Cartoon, topology of monomer. Representative of n=3 trials **(h)** SEC chromatogram of cleaved 2-pass LNLLx610 variant refolded and run in 16 mM C8E4 mobile phase; Inset, SDS-PAGE gel showing monomers (lower band) and SDS-resistant oligomers (upper band, assigned as pentamers) of 2-pass subunits. Cartoons, protein topologies. Representative of n=2 trials

**Supplementary Figure 7.**
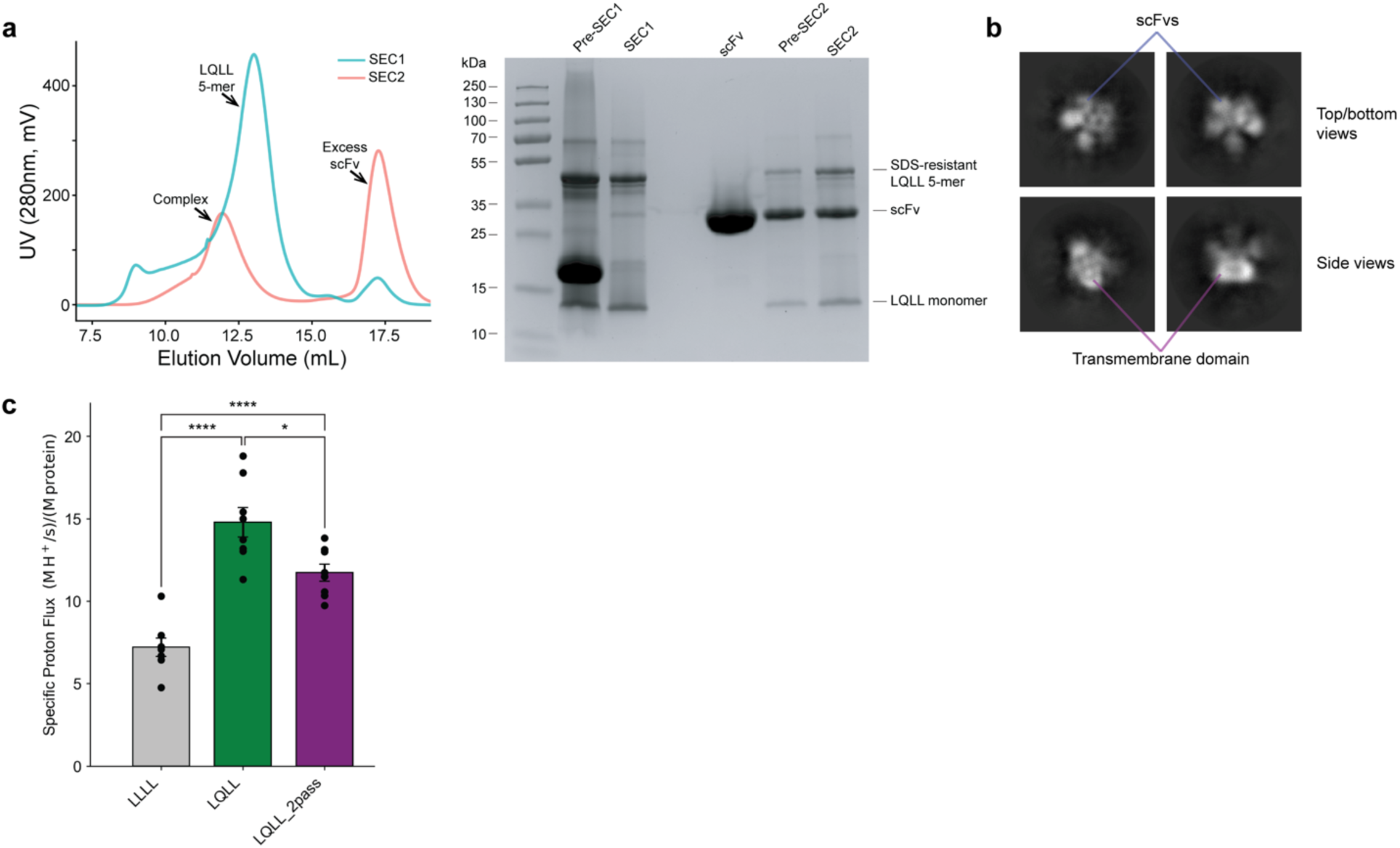
Pentameric scFv complex with LQLLx610-pA and relative liposomal flux. (**a**) Left, SEC chromatogram (UV, 280 nm) of 2-pass LQLLx610-protA at 1 mg/mL purified from Ni-NTA in C14B and exchanged into 0.05% DDM during SEC1 (blue trace). Sample was further complex with the 1.5 molar equivalent of 1P4B anti-helix antibody and run on s200i column with 0.05% DDM mobile phase (SEC2, red trace). Right, SDS-PAGE of co-eluting complex (cleaved sumo fusion at ∼16 kDa in pre-SEC1). Representative of n=3 trials **(b)** CryoEM 2D classes of scFv:LQLLx610-protA in DDM revealing classes of apparent pentamers, given the stoichiometry and arrangement of likely soluble domains **(c)** Relative proton conduction rates observed for 1-pass LQLL, 2-pass LQLLx610-protA, and non-conducting control variant 1-pass LLLL in liposomal flux assay (n=9 from N=3 independent trials), normalized by each protein’s relative reconstitution efficiency into liposomes measured by analytical HPLC.

**Supplementary Figure 8.**
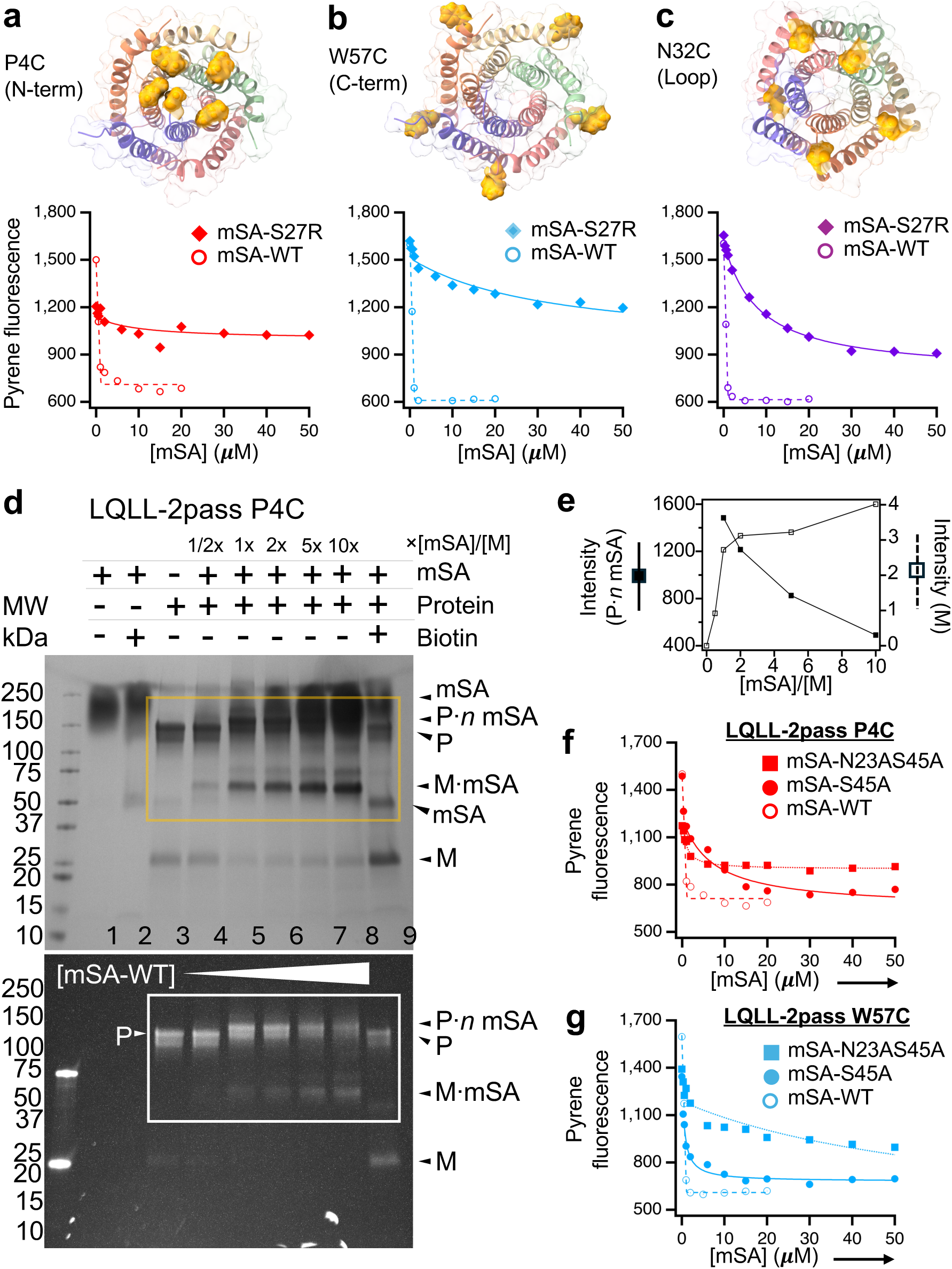
Evaluation of biotinylation conditions for steric trapping. (**a-c**) Identifying an optimal biotinylation site for steric trapping. All binding isotherms were obtained by monitoring quenching of pyrene fluorescence on the SUMO-fused LQLL proteins by dabcyl-labeled mSA. For each LQLL construct, an isotherm with strong biotin-binding mSA-WT was obtained to define the saturation level of fluorescence quenching. Three single Cys variants of LQLL-2pass for conjugating BtnPyr were evaluated in DDM. **(a)** P4C containing a Cys in the inner N-terminal helix. **(b)** W57C containing a Cys in the outer helix. **(c)** N32C containing a Cys in the loop connecting the inner and outer helices; and each variant was titrated with the weak biotin-binding variant mSA-S27R (*K*_d,biotin_ = 10^−5^ M)^42^. The P4C variant exhibited only the first binding phase with a negligible second binding phase, demonstrating that effective steric hindrance occurs between pre-bound and incoming mSA molecules. P4C was therefore selected as an optimal biotinylation site for steric trapping. In contrast, N32C showed a partial second mSA-binding phase, and W57C displayed a single, near-saturated binding phase. Both indicate that the biotin labels within the pentamer are not sufficiently close to generate effective steric hindrance between mSA molecules. **(d)** SDS–PAGE mobility shift assay demonstrating mSA-induced dissociation of TM domain pentamer. Dissociation of LQLL-2pass P4C pentamers (5 μM monomer equivalent) was monitored by adding increasing concentrations of mSA-WT from 0 to 50 μM (lanes 3–8). This assay exploits the strong, SDS-resistant mSA-WT– biotin interaction under non-boiled conditions to monitor mSA-induced dissociation of the pentamer^38,39^. As the mSA concentration increased, the intensity of the pentamer band (“P·*n* mSA”; migrating between 150–250 kDa) decreased, accompanied by a corresponding increase in the monomer band (M·mSA). Protein bands were assigned by comparing migration patterns of control samples (purified mSA: lanes 1–2; purified LQLL proteins: lane 3) and by contrasting Coomassie-stained gel (*top*, detecting all proteins) with UV-irradiated gel (*bottom*, detecting BtnPyr-labeled LQLL only). “P·*n* mSA” denotes the pentamer bound with *n* mSA molecules (*n* = 2 to 3 as shown in **Fig. 4a–c**). A band corresponding to the LQLL monomer bound to a single mSA molecule (“M·mSA”; migrating between the 50 and 75 kDa markers) was observed in the presence of excess mSA. **(e)** Band intensities from (d) for “P·*n* mSA” and “M·mSA” were quantified using ImageJ. **(f,g)** Selecting an mSA variant that optimally attenuates the second binding phase for steric trapping. The LQLL-2pass variants (P4C and W57C) were titrated with mSA variants of differing biotin affinities. For LQLL-2pass P4C **(f)**, the weaker biotin-binding mSA-N23A/S45A (*K*_d,biotin_ = 2 × 10⁻⁷ M)^42^ produces only a first binding phase, whereas the stronger biotin-binding mSA-S45A (*K*_d,biotin_ = 9 × 10⁻⁹ M)^38,42^ yields an optimally attenuated second binding phase resulting in full saturation of biotin sites. These results indicate that the combination of LQLL-2pass P4C and mSA-S45A provides effective coupling between mSA binding and pentamer dissociation, enabling reliable determination of pentamer stability.

**Supplementary Figure 9.**
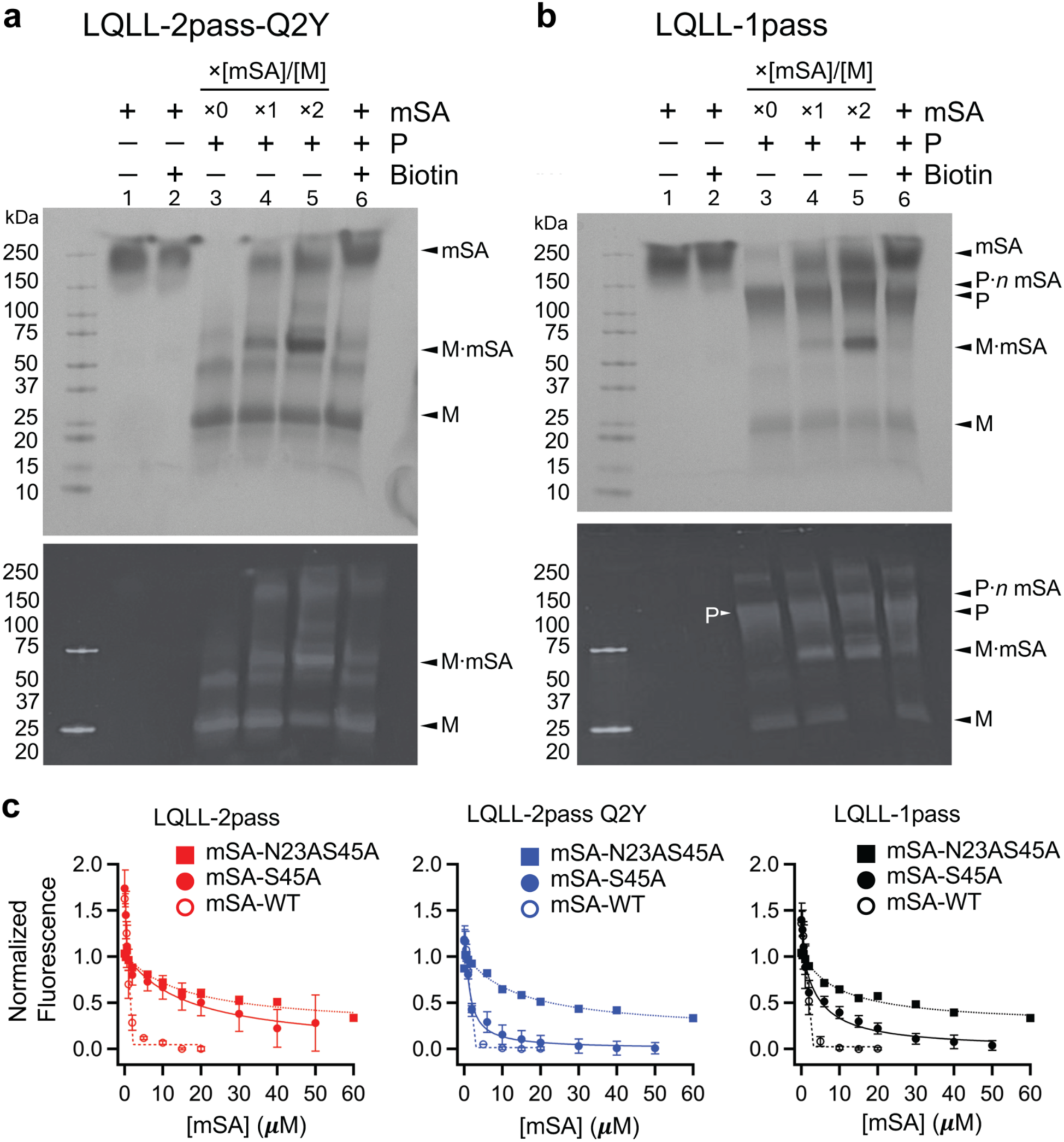
Steric trapping comparison of LQLL variants. (**a, b**) SDS-PAGE mobility shift assay verifying dissociation of LQLL variants via steric trapping. Two BtnPyr-labeled LQLL variants with an N-terminal SUMO fusion (MW: 21 kD; 5 μM monomer equivalent), **(a)** LQLL-Q2Y-2pass, and **(b)** LQLL-1pass, were incubated with increasing concentrations of mSA-WT (lanes 3–5; 0 μM, 50 μM, and 50 μM, respectively). Protein bands were assigned and labelled as in Figure S8d. Pronounced intensity in the LQLL-Q2Y-2pass sample in panel **(b)**, which strongly favors monomer formation. **(c)** Binding isotherms of mSA-S45A, mSA-N23A/S45A, and mSA-WT with 3 LQLL P4C constructs (left, LQLL-2pass; middle, LQLL-2pass Q2Y; right, LQLL-1pass). The Q2Y mutation destabilizes the pentamer interface, facilitating dissociation. In all cases, mSA-S45A produced appropriately attenuated second binding phases, confirming that it is optimal for steric-trapping measurements across all LQLL variants.

**Supplementary Figure 10.**
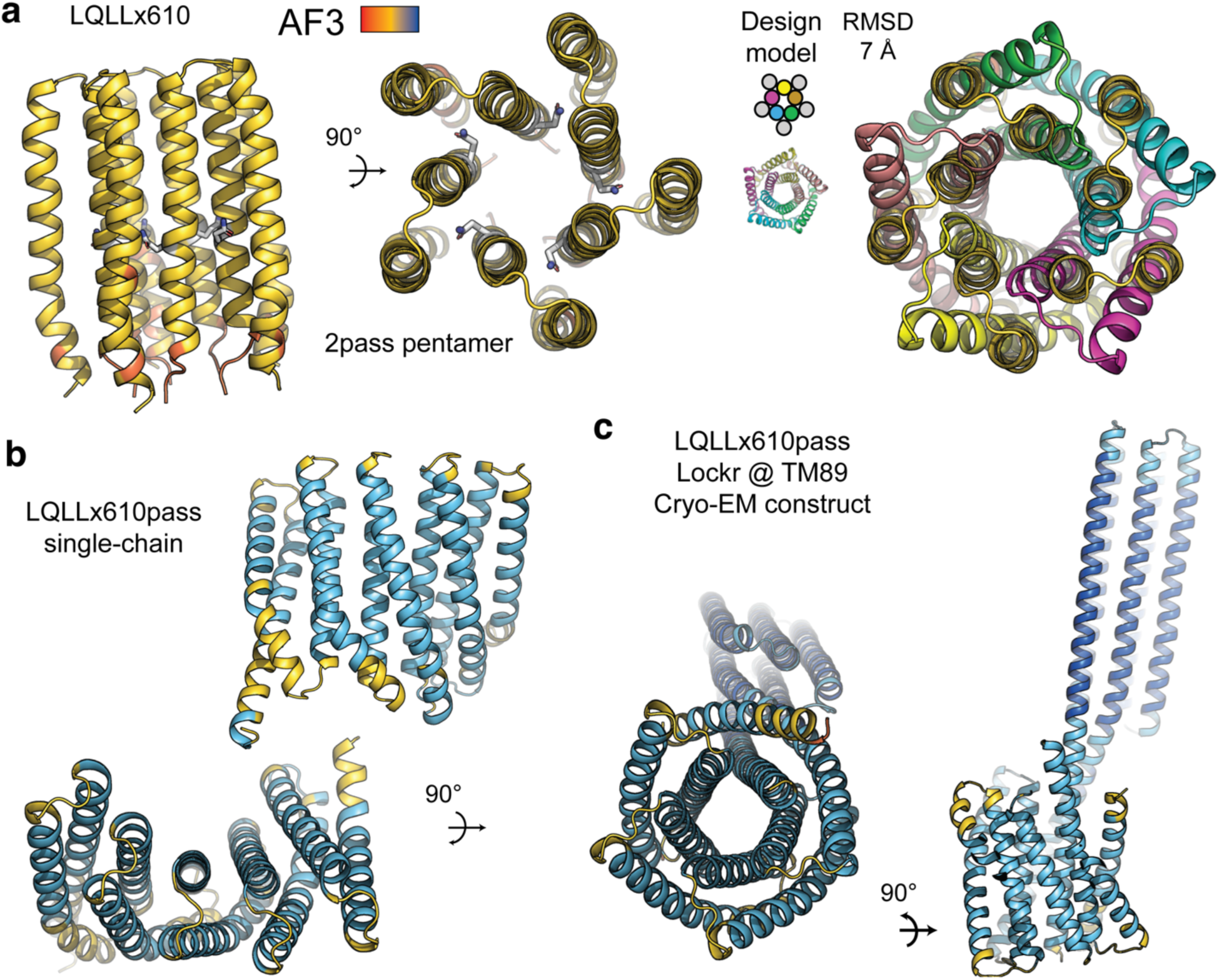
AF3 prediction of LQLLx610 2-pass pentamer and LQLLx610pass. (**a**) Left, AF3 prediction of LQLLx610 2-pass pentamer, colored by pLDDT, ipTM: 0.21; pTM: 0.29. Right, overlay with expected design model. **(b)** AF3 prediction of original LQLLx610pass design results in a designed helical bundle repeat-like fold (DHR)^57^, rather than the intended pseudo-symmetric channel structure, colored by pLDDT, pTM: 0.7. Redesign of the TM4-5 loop (variant LQLLx610*) converts the AF3 prediction, shown in in Figure 5. **(c)** AF3 prediction of LQLLx610pass Lockr @ TM89, cryo-EM construct, colored by pLDDT; pTM: 0.77.

**Supplementary Figure 11.**
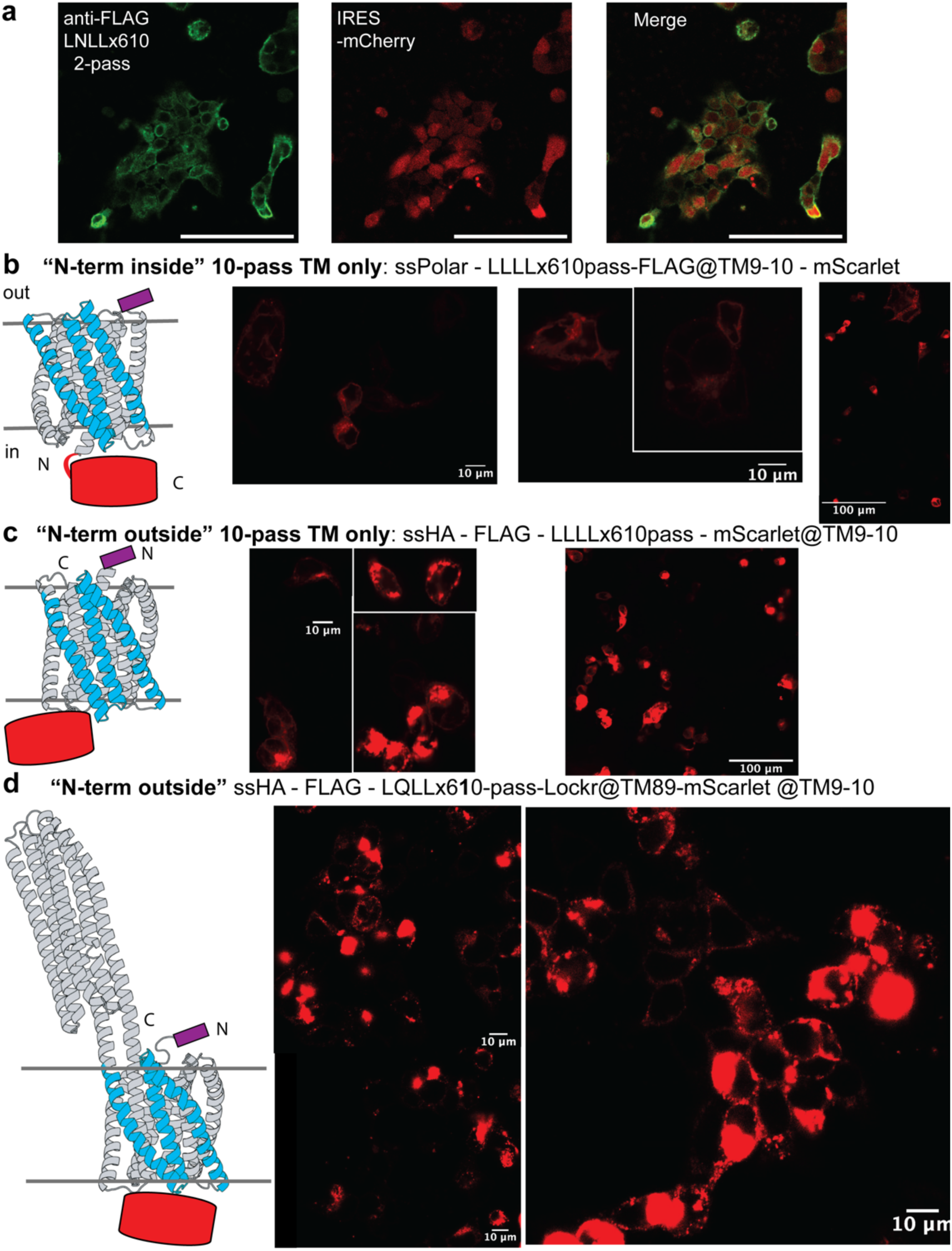
Membrane trafficking and topology control of multi-pass LQLLx610 variants. (**a**) Immunofluorescence staining of ssHA-FLAG-LNLLx610 transiently expressed in HEK293T cells (n=1). Left, anti-FLAG of non-permeabilize cells indicate membrane surfaced localized protein. Middle, mCherry expressed with cytoplasmic distribution from bicistronic element. Right, overlay. Scale bars, 100 µm. **(b)** Left, cartoon of LQLLx610pass construct with expected intracellular N-terminus, intracellular FLAG tag (purple), and extracellular mScarlet (red). Right, mScarlet fluorescence upon transient transfection shows plasma membrane cell surface localization (n=1). **(c)** Left, cartoon of LQLLx610pass construct with expected extracellular N-terminus and FLAG tag with intracellular cellular mScarlet. Right, mScarlet fluorescence upon transient transfection shows strong intracellular accumulation alongside membrane surface expression (n=1). **(d)** Left, cartoon of LQLLx610pass Lockr@TM89 construct with expected extracellular N-terminus and FLAG tag with intracellular cellular mScarlet. Right, mScarlet fluorescence upon transient transfection shows strong intracellular accumulation alongside membrane surface expression (n=1).

**Supplementary Figure 12.**
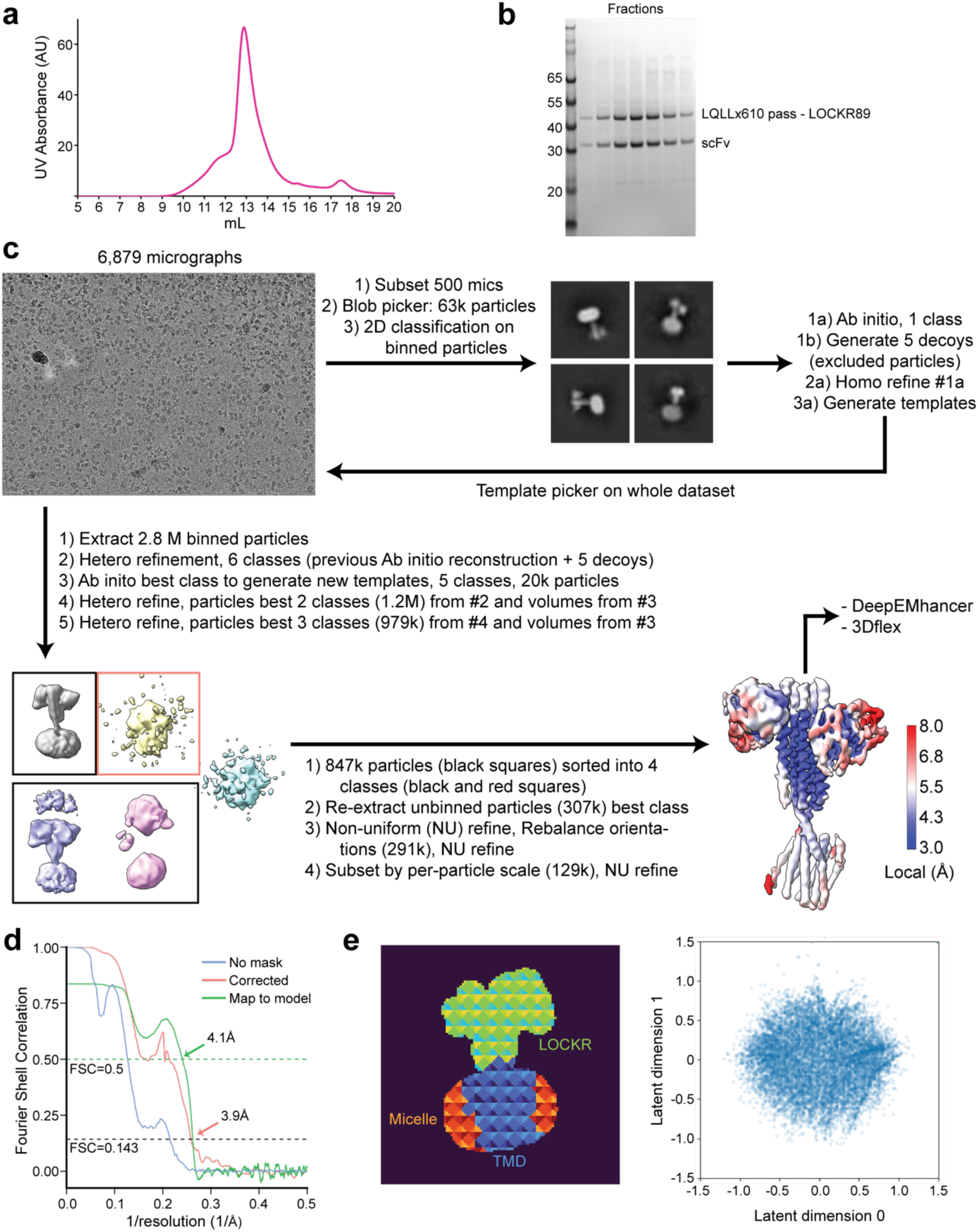
Cryo-EM Sample and Processing. (**a**) Size exclusion chromatography UV trace of LQLLx610pass-Lockr89:scFv complex and **(b)** corresponding SDS-PAGE of purified complex used for cryo-EM, both representative of n=3 independent trials. **(c)** Data processing flowchart for LQLLx610pass-LOCKR fusion. Output map for last NU refinement shown and colored by local resolution. This map was used for DeepEMhancer to produce final map, and as canonical density for 3DFlex analysis. **(d)** FSC curves for last NU refinement (no mask and corrected maps) and map to model. **(e)** Left, 3DFlex mesh used to separate protein into 3 domains: LOCKR+scFvs, TMD, and micelle (set as rigid). Right, scatter plot showing distribution of particle latent coordinates along the two latent dimensions.

**Supplementary Figure 13.**
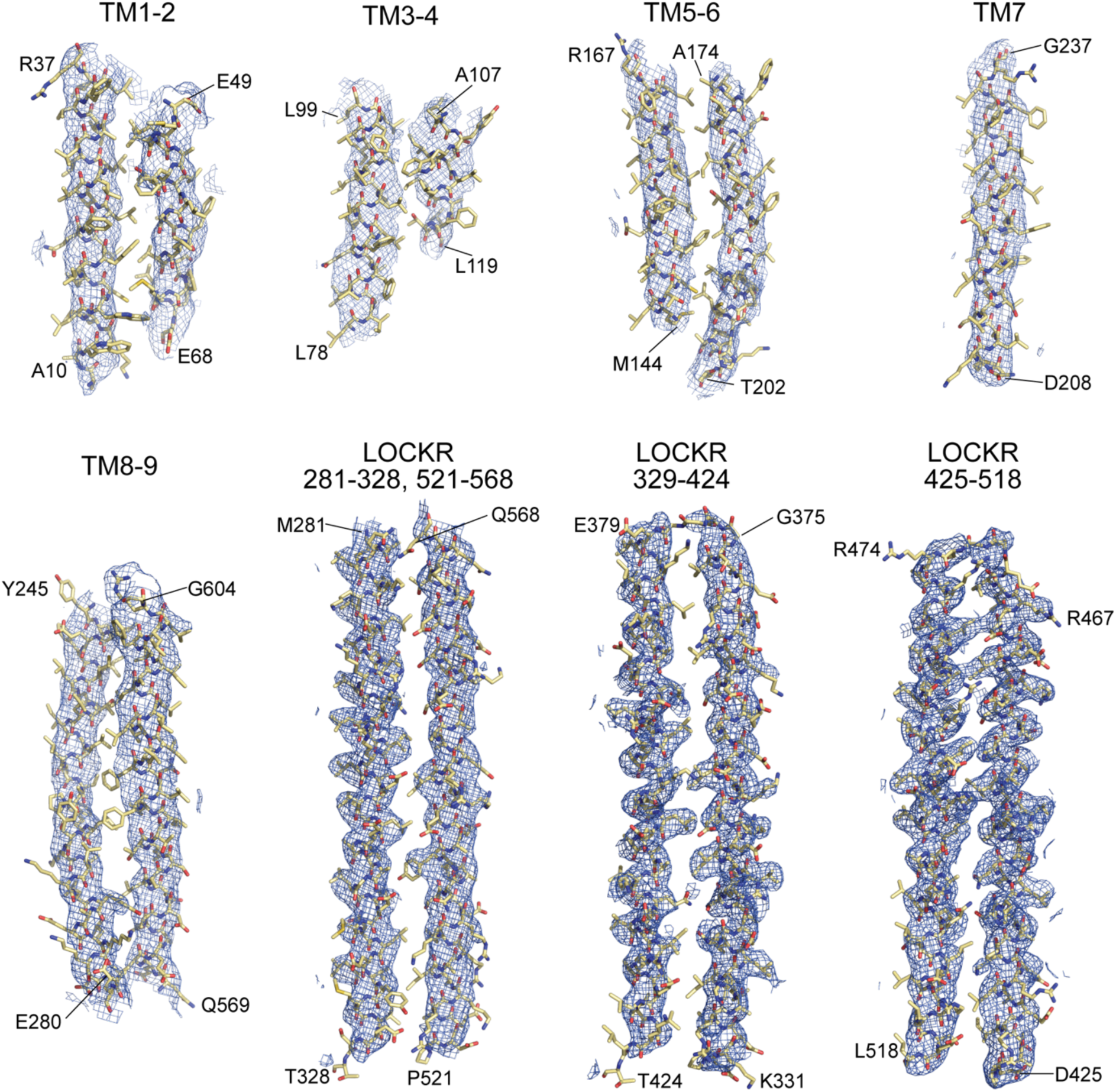
CryoEM density and fit to model of LQLLx610pass-Lockr89. Selected regions of atomic model of LQLLx610pass-Lockr89 superimposed onto its cryoEM map.

**Supplementary Figure 14.**
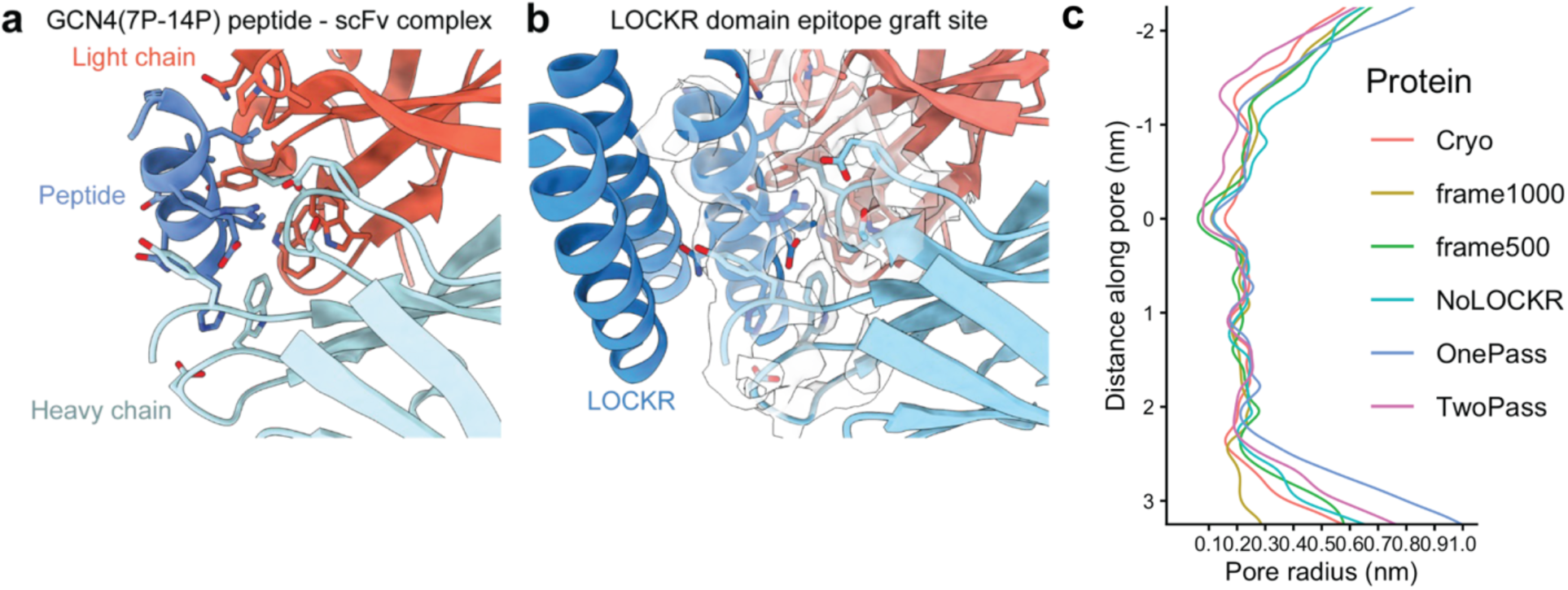
Selected interactions in LQLLx610pass-Lockr89 structure. (**a**) Depiction of interface between a helical peptide and its scFv (PDB:1P4B). **(b)** Binding interface of helical epitope 1P4B grafted onto LOCKR domain and its corresponding scFv. CryoEM map for LQLLx610pass-Lockr89 around interface is shown as density, confirming the conservation of the binding interactions shown in **(a)**. **(c)** Pore profiles of (red) LQLLx610pass-Lockr89 protein model fit into the cryo-EM density, (cyan) design model of LQLLx610pass, (blue) design model of LQLLx610, and (magenta) crystal structure of LQLL 1-pass overlaid with those of MD simulation frames starting from the model fit to cryo-EM density: gold, protein after 500 ns; green, protein after 1000 ns. MD simulation of LQLLx610pass-Lockr89 shows that its Q-ring pore constriction size (z = 0 nm) is similar to that of design models and the 1-pass LQLL experimental structure with modest fluctuations, consistent with the selective proton conduction mechanism.

**Supplementary Table 1.**
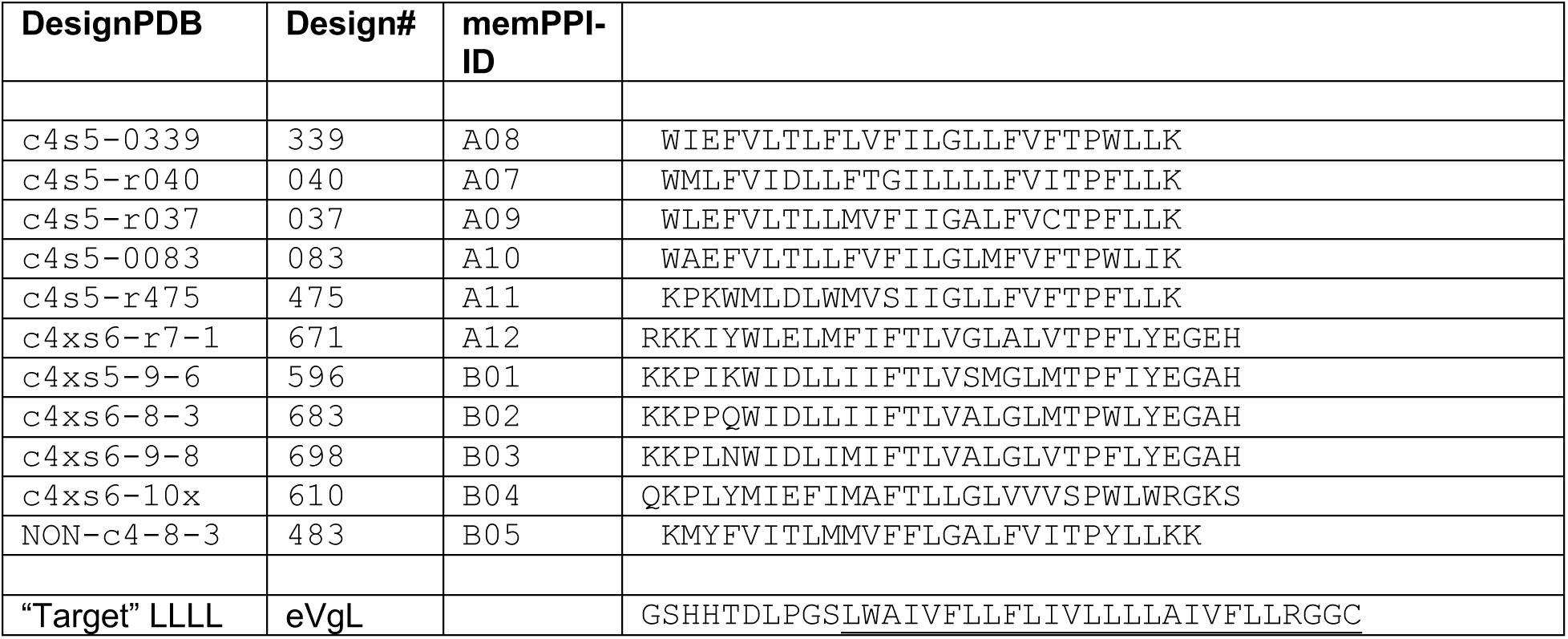
De novo designed TM adaptor domain sequences and expression constructs. Sequences derived from protein design models (and for *e. coli* protein expression).

**Supplementary Table 2.**
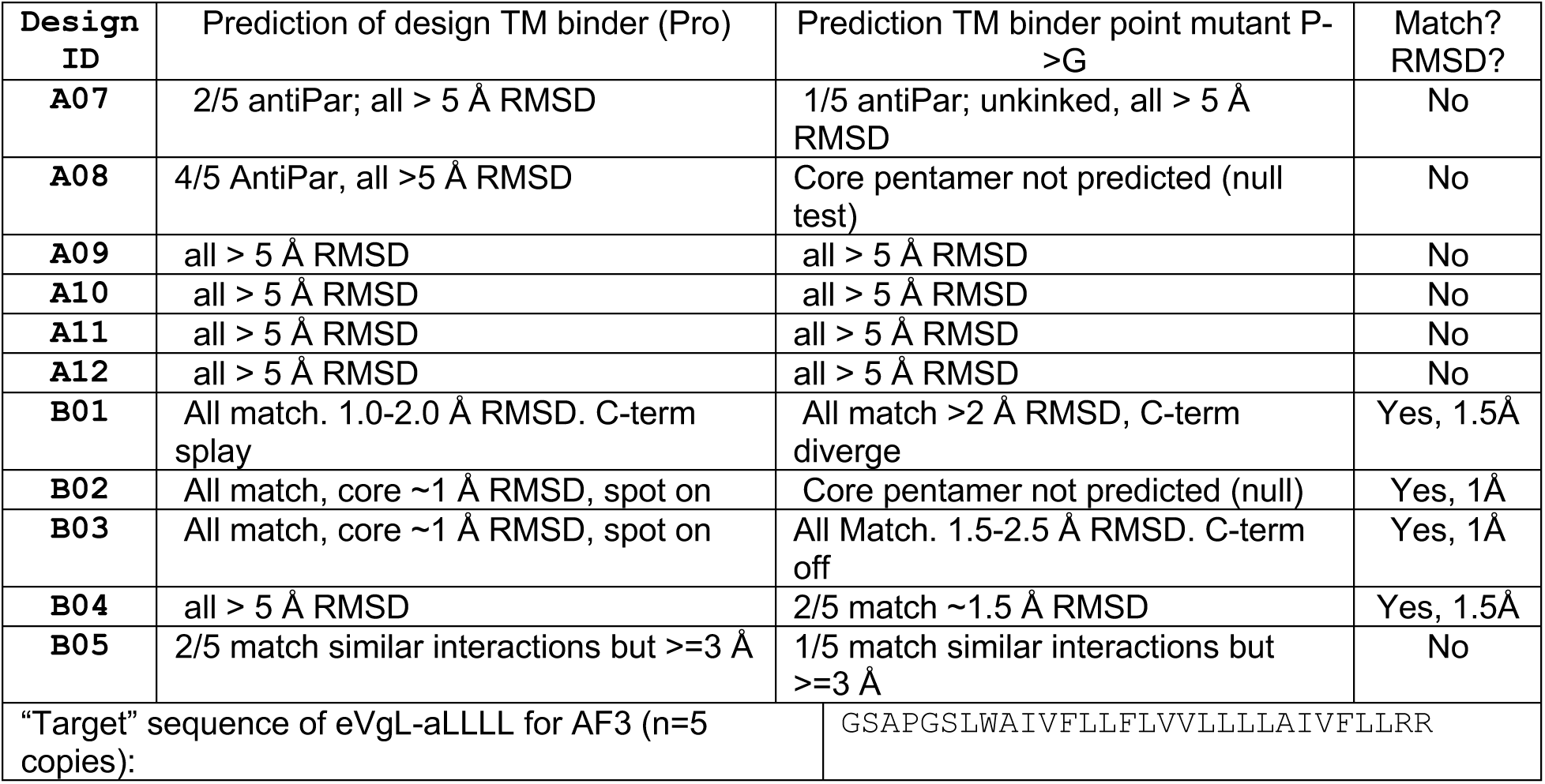
Summary of AlphaFold3 predictions of single-span TM adaptors co-complex with eVgL pentamer. Sequences derived from protein design models in Supplementary Table 1, intended for *e. coli* protein expression.

**Supplementary Table 3.**
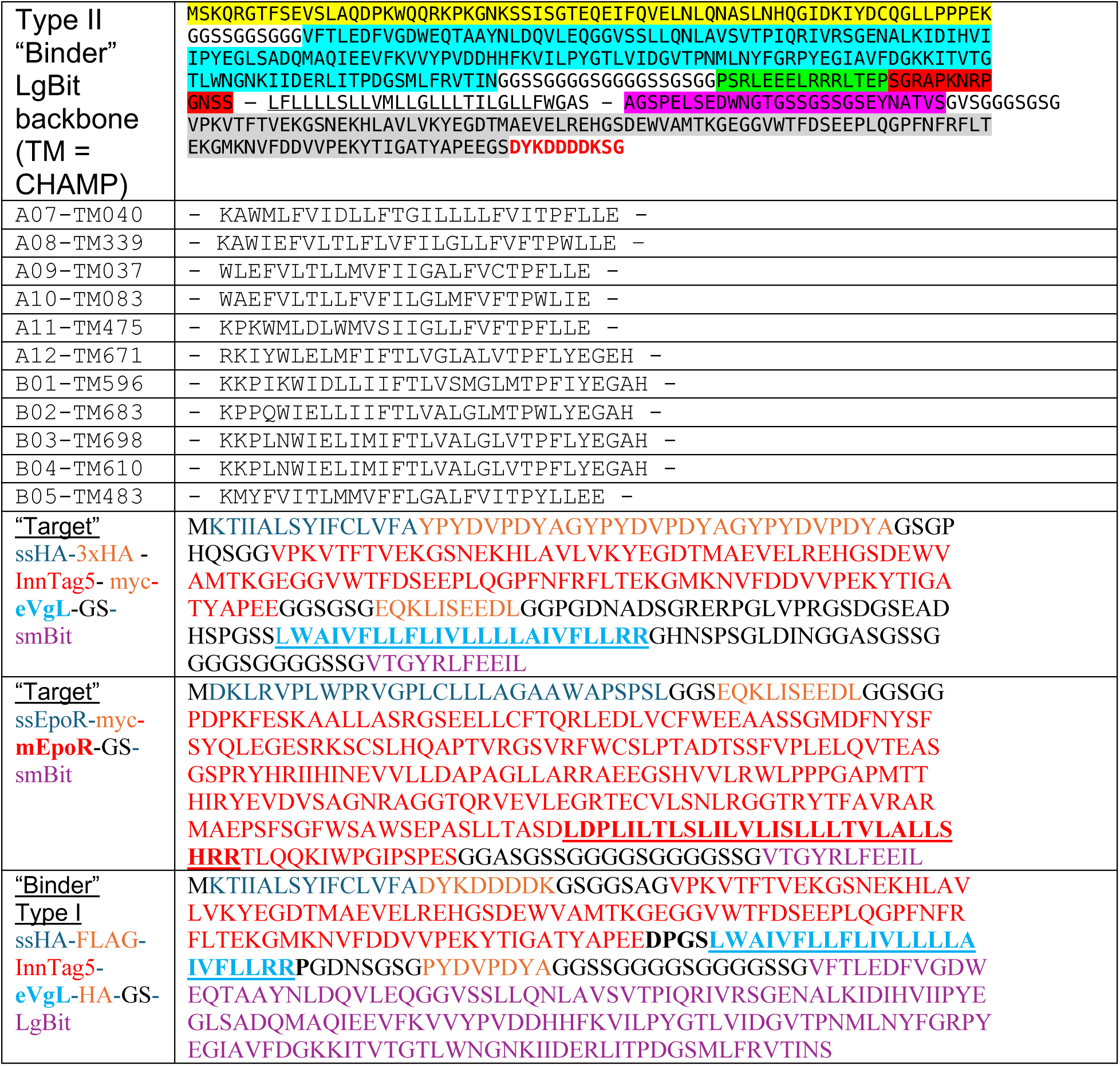
Protein sequences of expression constructs in mammalian cells for membrane protein-protein interaction (memPPI) NanoBit assay. Expression constructs in pcDNA3.1 transiently co-transfected for split nanoluciferase assay. Natural and synthetic “Target” SmBit-fusion proteins (Type I orientation) and “Binder” LgBit-fusion proteins (de novo designs intended Type II orientation). Expressed proteins’ plasma membrane surface trafficking and preferred insertion topologies were previously characterized in reference 33.

**Supplementary Table 4.**
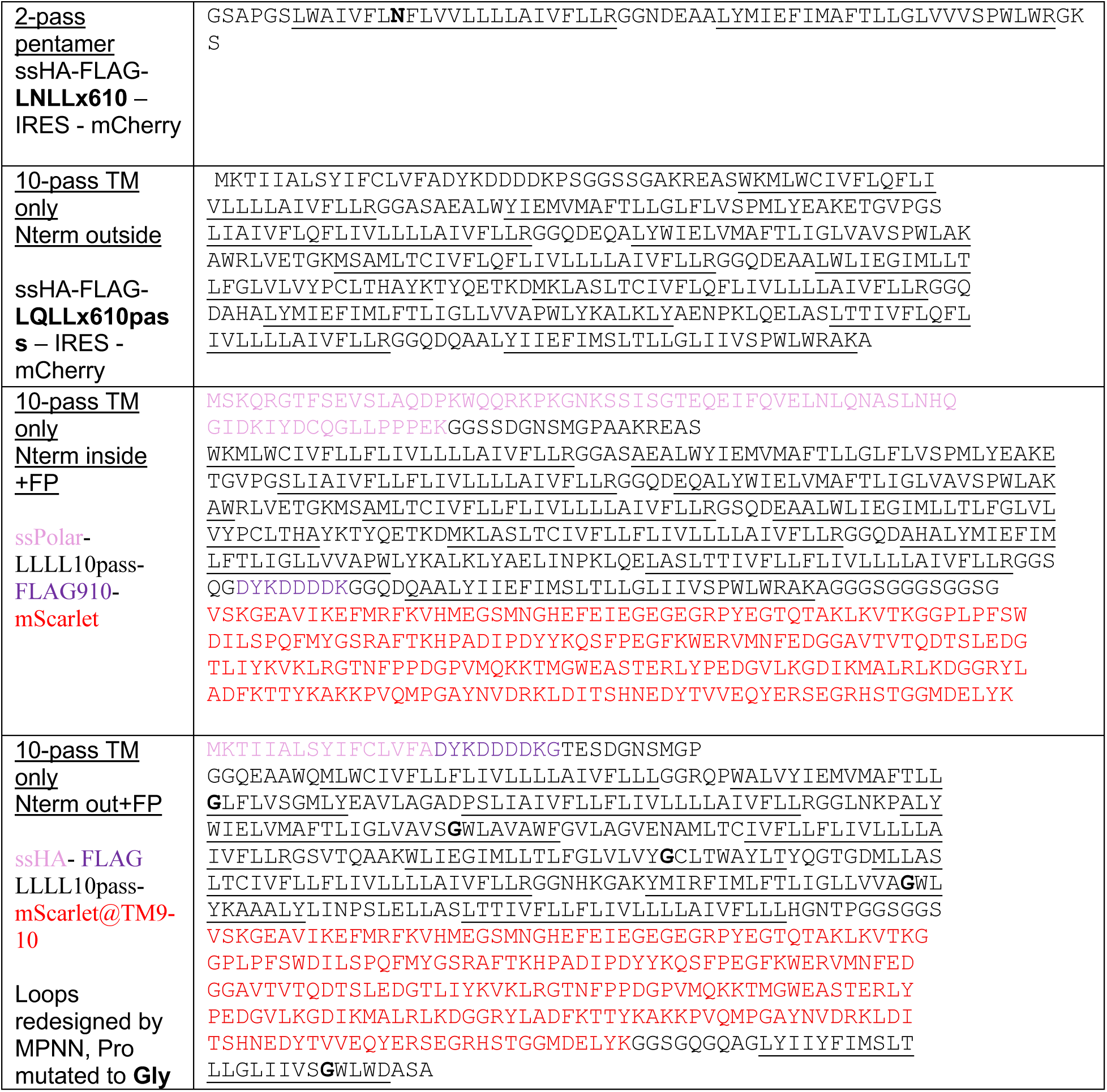
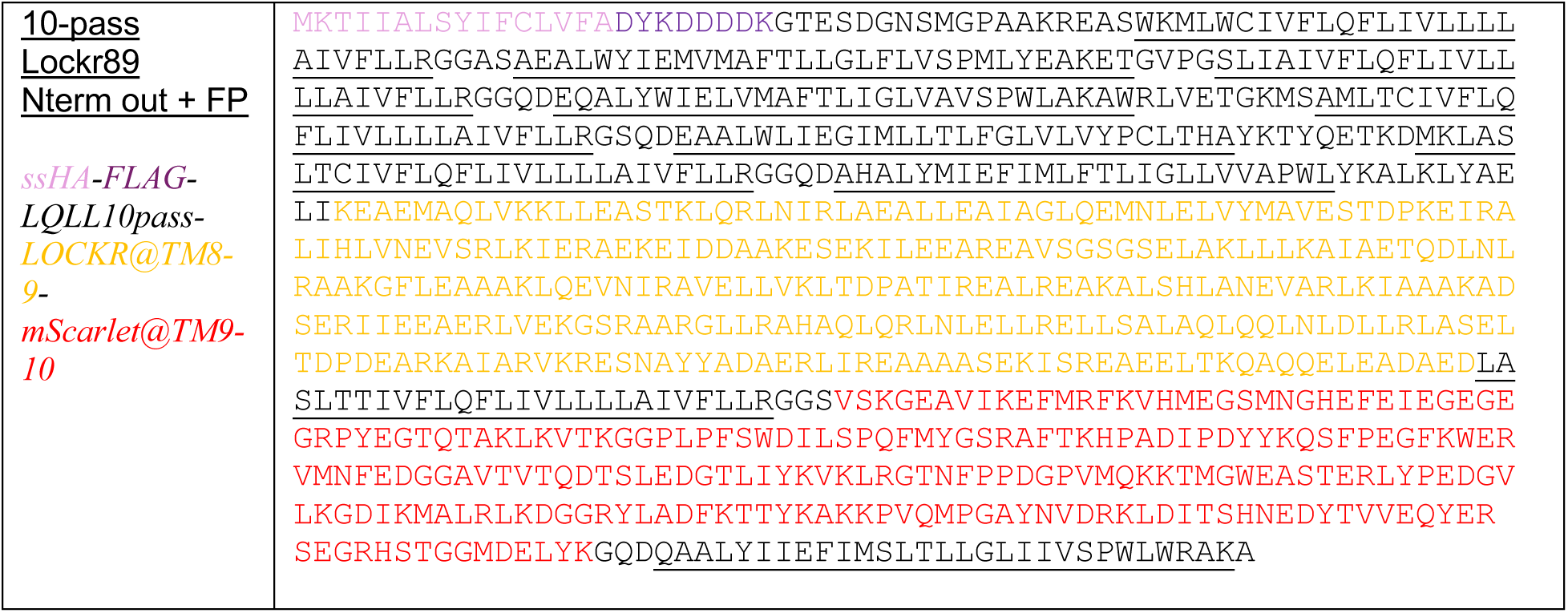
Protein sequences expressed in mammalian cells for membrane trafficking tests by fluorescence microscopy. Synthetic variants of multi-pass LXLL-610 proteins expressed by transient transfection. Transmembrane regions are underlined. Other domains such as fluorescent protein (FP) or Lockr@TM8-9 and other key sequence features (epitope tags; signal leader sequence: ssHA, ssPolar) are colored. Construct labels include as “out” or “in” tags to denote the expected extracellular or intracellular termini of the protein topology after plasma membrane trafficking.

**Supplementary Table 5.**
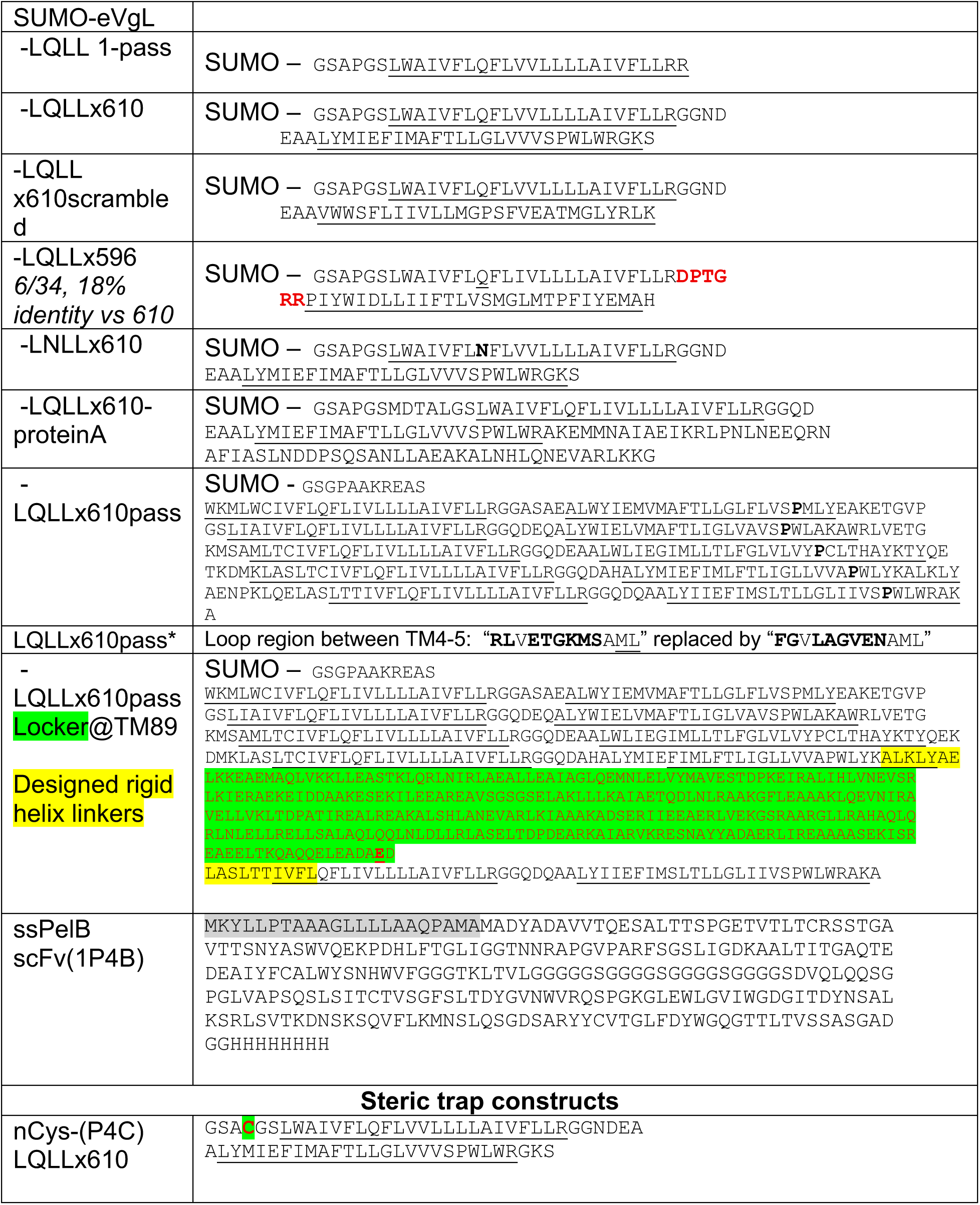
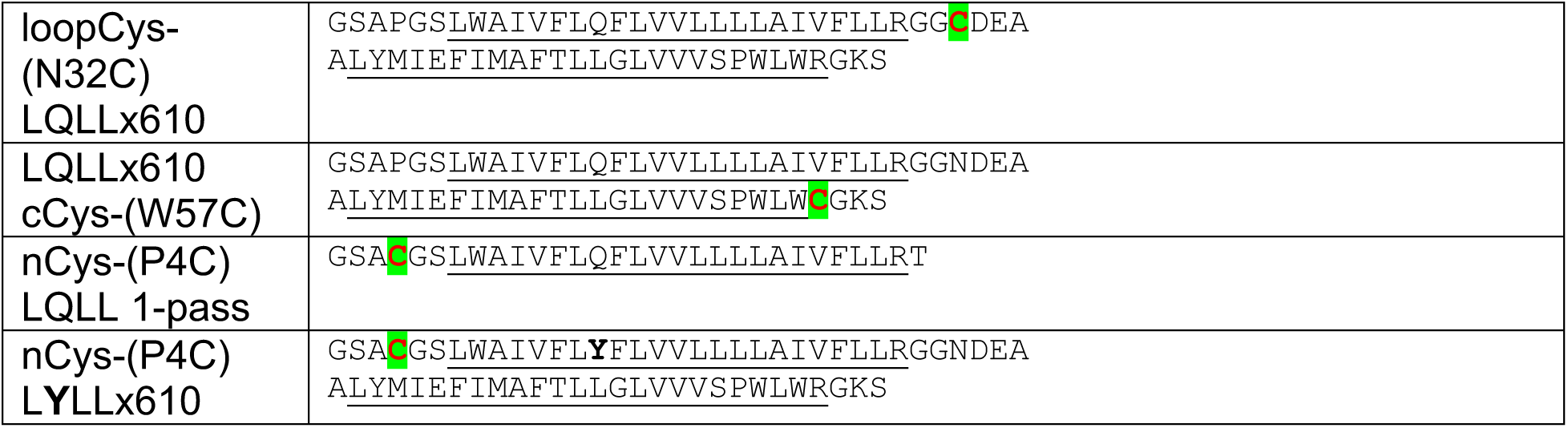
Protein sequences of bacterial expression constructs. His-tagged protein sequences expressed and purified from pET28 vector in C43(de3) *e. coli*.

